# Sialylated flagellin from *Pseudomonas aeruginosa* modulates the process of dendritic cell maturation through Siglec-9 mediated suppression of TLR5 signaling

**DOI:** 10.1101/2025.03.21.644494

**Authors:** Kaustuv Mukherjee

**Affiliations:** Cancer Biology & Inflammatory Disorder Division, CSIR-Indian Institute of Chemical Biology, Kolkata, India

**Keywords:** *Pseudomonas aeruginosa*, sialic acids, sialic acid-binding immunoglobulin-like lectins (siglecs), Siglec-9, dendritic cells (DCs), DC maturation, T cells, Toll-like receptor (TLR5)

## Abstract

Sialic acids, a nine carbon acidic sugar, can be detected in *Pseudomonas aeruginosa* (PA) when it is cultured with any exogenous sialic acids source like serum. Subsequently, several sialoglycoproteins from PA have been identified which includes flagellin. In the current study, the sialylation status of PA flagellin has been further investigated through lectin and Siglecs binding. Siglecs are immunoglobulin-like receptors which can recognize and bind to sialic acids. The sialylated PA flagellin (Fla^+Sia^) was observed to bind with its cognate receptor TLR5 (Toll-like receptor 5) and also with Siglec-9 on dendritic cell (DC) surface. DCs are potent antigen-presenting phagocytic immune cells which can activate other immune cells through costimulation. Immature DCs undergo maturation upon encountering antigens; which involves changes in the cell morphology, receptors expression, cytokine secretion etc. Here, immature DCs stimulated with Fla^+Sia^ or desialylated-Fla^+Sia^ (ds-Fla^+Sia^) underwent maturation but exhibited different phenotypic and functional profiles. The Fla^+Sia^ matured DCs showed relatively lower expression of costimulatory molecules, lower pro-inflammatory cytokines secretion, higher phagocytic activity and induced the generation of more Th (T helper) and Treg (regulatory T) cells. The impact of these two antagonistic signaling pathways-TLR5-based immune-activatory and immunosupressive Siglec-9 signaling on DC maturation was further explored in DCs generated from THP-1 cell line. Here, Fla^+Sia^ stimulated DCs showed increased Siglec-9-SHP1 association while ds-Fla^-Sia^ stimulation showed an increased TLR5 association with its interacting partners-MyD88/TRAF-6/IRAK-4. The TLR5/Siglec-9 signaling was disrupted through siRNA transfections or chemical inhibitor treatment which revealed that flagellin-based TLR5 stimulation was crucial for the initiation of robust cell signaling events. However, Fla^+Sia^ treatment activated Siglec-9 signaling, which directly inhibited TLR5 pathway. Interestingly, downregulating Siglec-9 signaling resulted in enhanced TLR5 activation upon Fla^+Sia^ stimulation, along with increased MAP kinase signaling and activation of the NF-κB pathway. Thus, Siglec-9 silenced DCs responded to Fla^+Sia^ with increased secretion of pro-inflammatory cytokines. Therefore, the altered profile and T cell polarization ability of Fla^+Sia^ matured DCs is caused by the Siglec-9-based disruption of TLR5 signaling.

## Introduction

*Pseudomonas aeruginosa* (PA) is a ubiquitous Gram negative bacteria often isolated from nosocomial infections (1, 2). Sialic acids are a family of nine-carbon containing acidic monosaccharides which are commonly found in higher vertebrates, but also in a few bacteria, protozooans etc (3). Sialic acid has been detected in PA when the bacteria are cultured in media supplemented with an exogenous sialic acid source (like serum or neuraminic acid, Neu5Ac) (4–6). Several PA proteins have also been identified to contain sialic acids, like the PA flagellin (6). Flagellin is the monomeric unit of flagella, the bacterial locomotory organ. Highly motile PA contains single polar flagella which aids in its virulence and dissemination (7). Based on reactions with antisera, PA strains express either a-or b-type flagellin. These two serotypes differ in the protein-coding sequences and are differently glycosylated (8–10). The flagellins of several Gram-negative bacteria are known to be glycosylated (11, 12). Flagellin glycosylation may facilitate bacterial virulence or prevent the detection of flagellin. Like, flagellin glycosylation enhances *Pseudomonas syringae* virulence against the host tobacco plants (13). In the case of *Burkholderia cepacia* (14) and *Campylobacter jejuni* (15), flagellar glycosylation may also dampen the host immune response against the highly immunogenic flagellin. As a highly conserved pathogen associated molecular pattern (PAMP), flagellins are easily detected by the host surface Toll like receptor-5 (TLR5). TLR-5-flagellin recognition leads to robust cell activation and the initiation of inflammatory responses (16).

Pseudaminic acid residues (a type of sialic acids) found in the flagellin of *Campylobacter jejuni*, allows it to interact with Siglecs in order to modulate the immune response against flagellin (16). Siglecs (sialic-acid-binding immunoglobulin-like lectins) are lectin-like receptors commonly found on haematopoietic cells (macrophages, dendritic cells, B cells etc) which can bind to sialic acids (17, 18). Many Siglecs play an additional role in modulating immune cell functions through signaling via cytoplasmic regulatory motifs. Some inhibitory Siglecs promote immunosuppression while a few activatory Siglecs are also known (17). Siglec-9 is a widely expressed inhibitory receptor containing ITIMs (immunoreceptor tyrosine based inhibitory motifs), which suppresses uncontrolled immune responses (19).

Dendritic cells are a class of highly phagocytic immune cells which function as efficient antigen-presenting cells (APCs) (20). DCs also express several costimulatory surface receptors (like CD80, CD86). Following antigen capture and processing, DCs can directly interact with and activate B and T lymphocytes, leading to their proliferation. Thus, DCs possess a remarkable ability to link both the innate and adaptive immune systems (20). Through its dual roles in antigen presentation and costimulatory activity, DCs can dictate the nature, type and vigor of T cell proliferation and responses against antigens. DCs are produced from the bone marrow CD34+ progenitor cells. These immature DCs may migrate to different tissues and reside there, remain in the bone marrow or remain free-floating in blood (21). Generally, immature DCs migrate to regions of inflammation or microbial attack where these cells encounter foreign antigens or chemical signals from the milieu, which induce immature DCs to undergo maturation. The lowering of endocytosis rate, the formation of dendrites and the increased expression of several costimulatory receptors are some of the hallmarks of DC maturation (20). Several types of DCs have been discovered to date and they vary in their surface markers and function. One method for studying DC activity involves stimulating blood monocytes to develop into monocyte-derived dendritic cells (moDCs) (22).

DC maturation can be triggered with microbial products like peptidoglycans, lipopolysaccharides (LPS) (23), lipoteichoic acid (24), flagellin (25) or genetic material like CpG DNA (26), viral mRNA, and dsRNA (27). Non-microbial but endogenous molecules and signals like cytokines (pro-inflammatory like TNF-α, IL-1β, type I IFNs), involvement of CD40L with CD40 receptor (20), HSP70, HSP90 (28) released into the microenvironment also causes DC maturation. DC surface Toll-like receptors (TLRs) are responsible for detecting several microbial products (29). The type of stimulus, as well as the involvement of TLRs plays an important role in determining the activity and response of matured DCs. Additionally, a variety of Siglecs are also expressed by DCs (30, 31). Generally, immune responses are regulated by the immune-inhibitory receptors like the Siglecs which prevent over-functioning of immune-activatory receptors like the TLRs. Interestingly, the involvement of inhibitory Siglecs during DC maturation influences its ability to stimulate T cells. This is particularly observed in tumor environments, where cancer-specific hyper-sialylated antigens promote the immune-suppression of DC functions through sialic-acid-Siglecs engagement (32, 33). While flagellin - a TLR5 ligand, has been used to induce DC maturation (25), we have investigated if the presence of sialic acids in PA flagellin has any impact on its role as an antigenic stimulus for the induction of DC maturation.

We have first established the presence of sialic acids in PA flagellin through biochemical assay, lectin and Siglec-binding assays. Such lectins, Siglecs binding are abolished after the enzymatic desialylation of flagellins (ds-Fla^+Sia^). We observed that Fla^+Sia^ binds to human monocyte derived dendritic cells (moDCs) through both TLR5 and Siglec-9. Next, we have explored the impact of flagellin sialylation on the process of DC maturation. Sialylated and desialylated PA flagellins were added to immature DCs and their maturation status was compared. A THP-1 cell line derived DC model was also used to dissect the interplay between TLR5 and Siglec-9 signaling pathways in greater detail. In this current study we report that that engagement of Siglec-9 by Fla^+Sia^ resulted in the activation of Siglec-based inhibitory signaling which led to the suppression of p38-MAPK, ERK1/2, Akt, NF-κB signaling cascades and also reduced the association of TLR5 with MyD88, IRAK4, and TRAF6. The inhibition of Siglec-9 signaling by siRNA based Siglec silencing or the inhibition of SHP-1/2 activity resulted in a greater activation of the TLR5 pathway in Fla^+Sia^ matured DCs. This leads to an increased secretion of pro-inflammatory cytokines. Together, these findings establish that the induction of Siglec-9 based inhibitory signaling by flagellin sialic acids interferes with TLR5 signaling. Such inhibitory signaling during the DC maturation process leads to altered DC phenotypes, which may ultimately favor the PA infection process.

## Materials and Methods

### Reagents

Roswell Park Memorial Institute 1640 (RPMI 1640), Fluorescein isothiocyanate (FITC), bovine serum albumin (BSA), 4’,6-diamidino-2-phenylindole (DAPI), Fluorescein tagged dextran beads (FD40S), paraformaldehyde, Anti-human IgG (Fc specific) antibodies peroxidase conjugated (A0170), TMB substrate (tetramethylbenzidine), Protein A-agarose beads (P9424), TH1020 (SML1741) and NSC-87877 (565851) were obtained from Sigma (St. Louis, MO). *Arthrobacter ureafaciens* sialidase (10269611001) was from Roche Applied Science (Mannheim, Germany). Vectashield mounting medium, biotinylated *Sambucus nigra* agglutinin (SNA) (#B-1305-2), biotinylated *Maackia amurensis* agglutinin (MALII) (#B-1265-1), agarose-SNA lectin beads (AL-1303-2) and biotinylated Peanut agglutinin (B-1075-5) were obtained from Vector Laboratories (USA). CFSE dye and PE conjugated anti-rabbit secondary antibody was from Molecular Probes, Thermo Fisher Scientific (OR, USA). Human TLR5 siRNA (sc-40262) and human Siglec-9 siRNA (sc-106550) was purchased from Santacruz Biotechnology (Texas, USA). CD83-PE (556855), CD209-FITC (551264), CD14-BV605 (564054), CD86-APC (555660), CD11c-BV650 (563404), CCR7-PE (552176), HLA-DR-APC (559866), IFN-γ-FITC (554700), TNF-α-FITC (552889), IL-12-APC (554576), IL-4-PE (554485), IL-10-PE (554498), CD3-PE (555340), CD3-V500 (561416), BD Tritest-CD3-PerCP, CD4-FITC, CD8-PE (340298), CD-25-PE (555432) and FoxP3 staining kit (560133) were obtained from BD pharmingen and BD Biosciences (San Jose, CA,USA).

Recombinant human Siglec-Fc chimeric proteins (Siglec-1, 3, 5, 7, 9), Recombinant Human GM-CSF (7954-GM-010), Recombinant Human IL-4 (6507-IL-010) were from R&D systems (MN, USA). All cell culture medium, fetal calf serum (FCS), human AB serum (H5667), lipofectamine LTX, plus reagent, Opti-MEM, cDNA synthesis kit, DyNamo Flash SYBR green etc. were purchased from Invitrogen (Thermo Fisher Scientific, Waltham, MA, USA). Pierce LAL Chromogenic Endotoxin Quantitation Kit, BCA assay kit, Firefly Luciferase Glow Assay Kit was purchased from Pierce (Thermo Scientific, Waltham, MA, USA). Clarity Western ECL Substrate, protein ladders, IPG strips, 2D clean up kits were obtained from BioRad (California, United States). All other antibodies were from Cell Signaling Technologies (MA, USA) unless indicated otherwise. Polyvinylidene difluoride membranes (PVDF) were obtained from Millipore (USA) and molecular weight cut off centrifugal filters (Vivaspin) from GE healthcare. Mass-spec grade Trypsin Gold (V5280) was obtained from Promega.

### Bacterial culture

*Pseudomonas aeruginosa* strain PA14 was a gift from Prof. Richard D. Cummings (Emory University School of Medicine, Atlanta, USA) (4). As established previously, non-sialylated PA (PA^−Sia^) is obtained by culturing this bacteria in only tryptic soy broth (TSB) at 37°C overnight under shaking condition (4–6, 34). Sialylated PA (PA^+Sia^) was obtained by culturing the bacteria overnight in TSB supplemented with 10% heat-inactivated FCS under the same conditions. *Escherichia coli* ATCC 25922 was obtained from Dr. Rupak Kr. Bhadra, CSIR-IICB, Kolkata and cultured in TSB.

### Cell culture

Human monocytic (THP-1) cell line was sourced from the National Cell Repository of the National Centre for Cell Science, Pune, India and routinely cultured in RPMI 1640 medium supplemented with 10% FCS and incubated at 37°C with 5% CO2. THP-1 derived dendritic cells (DCs) were prepared as per Berges C et al, 2005 by culturing THP-1 cells in RPMI with 10% FCS supplemented with rh-GMCSF (100 ng/mL) and rh-IL-4 (100 ng/mL) (35). Cells were cultured for 5 days with medium change after every 2 days of culture. Cell differentiation into DC like cells was confirmed through the detection of DC specific markers.

Human monocyte derived dendritic cells (hmoDCs) were generated from peripheral blood mononuclear cells (PBMCs) as per Nair S et al, 2012 (36). Blood was collected from 4 healthy donors after obtaining their informed consent in accordance with the Declaration of Helsinki after approval from The Ethical Committee on Human Subjects, CSIR-IICB, Kolkata. PBMCs were isolated from the collected blood through centrifugation using histopaque and then treated with ACK lysis buffer (Gibco). The PBMCs were then incubated in serum-free IMDM medium in T75 flasks for 2 hr at 37°C. Through plastic adherence, monocytes remained attached to the flask and all floating cells were removed. Monocytes isolated in this manner were further incubated in RPMI 1640 medium with 10% FCS supplemented with rh-GMCSF (50 ng/mL) and rh-IL-4 (50 ng/mL) for 5 days. Medium was refreshed every 2 days and cellular differentiation was monitored through changes in cell morphology as well as a loss of attachment. Expression of DC markers through flow cytometry confirmed the generation of immature moDCs.

These immature moDCs or THP-1 derived DCs were incubated with different doses of Fla^+Sia^ /ds-Fla^+Sia^ for 24 hours at 37°C under 5% CO_2_ to induce maturation. For studies involving cell signaling, THP-1 derived DCs cells were stimulated overnight with 2 µg of flagellin.

### Lectin based pulldown of sialoglycoproteins from PA^+Sia^

Overnight cultured PA^+Sia^ were lysed in PBS containing protease inhibitor by gentle sonication. This bacterial lysate (100 μg) was incubated with *Sambucus nigra* (SNA) lectin conjugated to agarose beads overnight at 4°C. Beads were gently washed with chilled PBS, pelleted by centrifugation and incubated in SDS sample buffer in order to extract all the bound proteins. Samples were resolved via 10% SDS-PAGE and the gel was stained with Coomassie Brilliant Blue 250 dye. Destaining revealed several fine bands, two distinct bands near 34.5, 37.5 kDa (from SNA lectin) and one prominent band near 47kDa. This prominent band was excised and then processed using in-gel tryptic digestion kit (Pierce, Thermo scientific).

### Isolation of flagellin

Flagellin was isolated from PA cultured with (PA^+Sia^) or without FCS (PA^-Sia^) following the method described by VL Campodónico et al, 2010 (37). PA was cultured overnight with gentle shaking at 37°C followed by pelleting by centrifugation at 5000 rcf for 10 min. PA pellet was resuspended in 20 mM Tris-Cl buffer (pH 7.4) and flagella was sheared by gentle homogenization using a french press. Intact bacterial cells were removed from the suspension by high speed centrigugation at 13,000 rcf for 30 min at 4°C. The supernatant was collected and flagellin was precipitated by salting out using 20% ammonium sulfate. After overnight incubation, the precipitate was extensively dialyzed against 20 mM Tris-Cl buffer to allow the protein pellet to redissolve. The purified flagellin proteins were then treated with agarose-bound polymyxin B sulfate beads to remove any contaminating endotoxins. Beads were added to the purified protein (1:3 v/v) and incubated for 2 hours in ice with intermittent mixing. The solution was centrifuged to pellet the beads and the supernatant containing flagellin was collected.

The absence of any contaminating endotoxins was always confirmed using the LAL Chromogenic Endotoxin Quantitation Kit (88282, Pierve, Thermo Scientific) before cell stimulation was performed. The protein concentration was estimated by BCA method and the purity was checked via SDS-PAGE (10% gel).

### Desialylation and FITC-labeling of flagellin

Purified Fla^+Sia^ (25 µg) was overnight incubated with *A. ureafaciens* sialidase (10mU/100μL) in acetate buffer (pH 5) at 37°C to obtain desialylated flagellin (ds-Fla^+Sia^). Proteins were resolved in SDS-PAGE to confirm that proteins were not degraded following incubation.

For FITC labelling, Fla^+Sia^ (100 μg) was incubated with 0.1% FITC in bicarbonate buffer (50 mM, pH 8.0) for 45 min at 37°C. The proteins along with the labeling solution were transferred to centrifugal filters with 15 kDa molecular weight cut off. All excess, unbound FITC which remained with the labeled protein were removed through extensive buffer exchange against 20 mM Tris-Cl buffer using this centrifugal filter. The presence of FITC tag in Fla^+Sia^ was confirmed by fluorimetry before experiments were performed.

### Acetyl acetone based fluorimetric estimation

Flagellins isolated from PA^-Sia^/PA^+Sia^(500 μg) were processed for acetyl acetone based detection of sialic acids as described by Shukla et al, 1982 (38). The proteins were treated with periodate solution (2.5 mM) at 4°C for 15 min in dark for mild oxidation followed by incubation with sodium arsenite (2% in 0.5M HCl) and acetyl acetone (750 mL glacial acetic acid, 3.75 g Ammonium acetate and 500 μL of acetyl acetone in 250 mL water) at 60°C for 10 min consecutively. The reaction was stopped by the addition of 2.5 mL of water. The sialic acids present in protein solution forms a fluorogenic product after the reaction which was detected by fluorimetry with excitation at 410 nm and emission at 510 nm wavelength. A standard curve prepared from known concentrations of commercially available N-acetyl neuraminic acid was used to quantify the sialic acid content.

### Isoelectric focussing (IEF) and 2D gel electrophoresis

Purified Fla^+Sia^ (25 µg) was precipitated by overnight incubation with chilled acetone at-20°C. These proteins were resolubilized in rehydration buffer (8 M urea, 2% CHAPS, 50 mM DTT, 0.2% Bio-Lyte 3/10 ampholyte, 0.001% Bromophenol Blue) and then passively rehydrated into IPG (immobilized pH gradient) gel strips (pH 4-7) as per manufacturer’s protocols. IEF was carried out in a 3 step program - 20 min at 250 V at linear mode; 2 h at linear mode from 250 to 4000 V; at 4000 V up to 10000 V-h in rapid mode in a Protean IEF Cell (Biorad).

Following isoelectric focusing, IPG strips were treated with equilibration buffer I (6 M Urea, 2% w/v SDS, 30% glycerol, 2% w/v dithiothreitol, 50mM Tris-HCl pH 6.8) for 30 min at room temperature followed by equilibration buffer II (6 M Urea, 2% w/v SDS, 30% glycerol, 2.5% w/v iodoacetamide, 50mM Tris-HCl pH 6.8) for 30 min. After reduction, alkylation, IPG strips were placed on top of 10% acrylamide gel using low temperature melting overlay agarose (1632111, BioRad) and then resolved in the second dimension by SDS-PAGE. The 2D gel was stained with Coomassie dye and destained to observe protein spots.

In a similar setup, Fla^+Sia^ and ds-Fla^+Sia^ (20 μg) was also rehydrated into IPG strips (pH 3-10) and resolved by isoelectric focusing following the same 3 step program. IPG strips were fixed, stained by Coomassie blue and destained to visualize the protein resolution.

### Mass spectrometry based identification of proteins

Coomassie stained protein bands were carefully excised and then destained (20mM ammonium bicarbonate in 50% acetonitrile). The gel pieces were processed using in-gel tryptic digestion kit (Pierce, Thermo scientific). The protein samples in bands were reduced (50 mM TCEP in 25 mM ammonium bicarbonate buffer, incubated at 60°C for 10 min), then alkylated (100 mM iodoacetamide in 25 mM ammonium bicarbonate buffer, incubated in the dark at room temperature for 1 hr) followed by washing with 50% acetonitrile. The gel pieces were then completely dehydrated by incubating with 100% acetonitrile for 15 mins and then dried in a laminar air flow hood. These gel pieces were then rehydrated with trypsin (100 ng/ band) in 25 mM ammonium bicarbonate buffer and incubated overnight at 37°C. The digested peptides were extracted from gel pieces using successive washes with acetonitrile (50%) and trifluoroacetic acid (0.1%) solution and pooled. The pooled digest was dried in a centrifugal evaporator. The dried peptide mix was dissolved with α-cyanohydroxycinnamic acid (CHCA) matrix before spotting on to MALDI-TOF/TOF plates. For calibration, spectra of calmix was used and samples were analyzed using MALDI mass spectrometer (ABI-4800 from Applied Biosystems, Foster City, CA). For the detection of peptide peaks, the scan range was fixed at 700-4000 (m/z). Both MS and MS/MS spectra were analyzed by MASCOT search engine version 3.5 based peptide mass fingerprinting using the ABI GPS Explorer software (Applied Biosystems). The protein identity was confirmed using BLAST tool based searching of eubacterial protein spectra database with following parameters - cleavage enzyme: trypsin; variable modifications: oxidation on methionine; fixed modification: carbamidomethylation of cysteine; missed cleavages permitted: one missed cleavages; minimum signal to noise ratio (S/N): 10; peptide charge: +1; precursor mass tolerance: ± 100 ppm; mass tolerance for the MS/MS search: ± 0.2 Da.

The purified flagellin band (2 µg) was similarly processed and subjected to trypsin digestion. The dried tryptic digest was reconstituted in 70% acetonitrile containing 0.1% formic acid. About 5µL of the digest was injected and first fractionated by a C18 easy spray nano column fitted in the nano-LC (Easy-nLC1000) using a gradient elution prepared from the mixing of mobile phase A (water and 0.1% formic acid) and mobile phase B (acetonitrile and 0.1% formic acid). The released samples were subjected to ESI-MS/MS in the positive mode using the LTQ-Orbitrap mass spectrometer (LTQ-XL, Thermo-Fisher Scientific). For the detection of peptide peaks, the scan range was fixed at 350-3000 (m/z). The resulting peak data were searched against the Uniprot eubacteria database using Sequest HT Proteome Discoverer 1.4 software (Thermo Fisher Scientific) utilizing the MASCOT search engine. The false discovery set at a cut off ≤1%. Following search parameters were set - cleavage enzyme: trypsin; variable modifications: oxidation on methionine; fixed modification: carbamidomethylation of cysteine; missed cleavages permitted: one missed cleavages; minimum signal to noise ratio (S/N): 10; peptide charge: +1; precursor mass tolerance: ± 100 ppm; mass tolerance for the MS/MS search: ± 0.2 Da

### Detection of the association between flagellin and Siglec-Fc by ELISA

Specifically 100 µL of Fla^+Sia^ in bicarbonate buffer (2µg/mL) was aliquoted into ELISA plates and incubated overnight at 4°C for coating. The coating solution was removed, the plates were washed with PBS and the wells were blocked with 4% desialylated BSA in Tris Buffered Saline (TBS) buffer for 1 hour. The blocking solution was removed; wells were incubated with 100 µL of recombinant Siglec-Fc chimeric proteins (Siglecs-3, 4, 5, 6, 7, 9 and 10-Fc, RnD systems) which were reconstituted in sterile PBS (20 µg/mL). Following 3 hours of incubation at room temperature, wells were again washed with PBS and 100 µL of anti-human IgG (Fc specific) peroxidase (1:20,000) in blocking solution was added. After 1 hour incubation at room temperature, wells were washed with PBS (5 min intervals, 6 times). The extent of Siglec-Fc bound to flagellin-coated wells was detected by incubating with TMB substrate. The reaction was stopped by the addition of 0.1 N HCl and the colored product formed due to peroxidase activity was measured at 450 nm by an ELISA reader.

### Lectin and Siglec blotting

Fla^+Sia^/Fla^-Sia^ (5μg) were resolved in 10% SDS-PAGE and transferred onto PVDF membranes by the wet transfer method. The blots were incubated with 2% desialylated-BSA in TBS buffer for blocking followed by incubation with biotinylated *Maackia amurensis* lectin II (MALII) or biotinylated *Sambucus nigra* lectin (SNA) (5 μg/mL) overnight at 4°C. Blots were washed using TBS-Tween 0.1% and probed using avidin-HRP (1:10,000). Any lectin binding was detected through reactions between HRP and chemiluminescent substrate and visualized in a chemidoc. The Ponceau S stained images of the blots have been used to account for equal protein loading.

In a similar manner, Fla^+Sia^ (5μg) resolved in 10% SDS-PAGE was transferred on to blots. These blots were incubated overnight at 4°C with Siglecs-3,4,5,7,9,10-Fc chimeric proteins (20 µg/mL) respectively. Blots were processed similarly to western blotting and the binding of different Siglec molecules with Fla^+Sia^ was detected using anti-human IgG (Fc specific)−peroxidase (1:5000). Equal loading of Fla^+Sia^ across all blots were confirmed from the Ponceau S staining of blots.

## Flow Cytometric Analysis

### Detection of Fla^+Sia^ binding with human monocyte derived DCs (hmoDCs)

Immature hmoDCs were incubated with anti-Siglec-5 (MAB1072, RnD Biosystems) or anti-Siglec-9 antibodies (MAB1072, RnD Biosystems) (1:1000) for 1 hour at 4°C. These cells were washed with PBS and then further incubated with phycoerythrin (PE) labeled anti-murine secondary antibodies for 30 min at room temperature. The cells were finally washed with PBS and then analyzed by flow cytometry.

Similarly, immature hmoDCs were treated with anti-Siglec-9 or anti-TLR5 antibodies (1:1000) for 1 hr at 4°C. Antibody-blocked and control hmoDCs (1×10^6^) were incubated with FITC-Fla^+Sia^ (2μg) at 4°C for 1 hour followed by PBS wash and flow cytometry. Cells were analyzed in FACS LSR Fortessa (BD Biosciences) flow cytometer and acquired data was analyzed in FACSDiva 8.0.2 software.

### Detection of surface markers and internal cytokine levels in human monocyte derived DCs (moDCs)

After 24 hour stimulation with Fla^+Sia^/ds-Fla^+Sia^ (1 μg, 2.5 μg, 5 μg), the matured hmoDCs were washed with PBS and incubated in PBS-BSA (0.5%). Fluorophore-conjugated antibodies against respective surface markers were added to these cells following manufacturer’s instructions and cells were incubated for 30 min at 4°C. The antibodies were combined into different cocktails as per the following combinations – (CD83-PE, CD86-APC, CD11c-BV650); (CCR7-PE, HLA-DR-APC). The unstained cells, cells stained with single-color antibodies and cells stained with antibody cocktail were pelleted, washed with PBS, and then analyzed in a flow cytometer. The single colour stained samples were used for compensation, when required. Data was acquired in a FACS LSR Fortessa (BD Biosciences) flow cytometer and then analyzed via FlowJo VX software.

For the detection of intracellular levels of cytokines, matured DCs were treated with fixation buffer (BD cytofix) followed by the permeabilization buffer (BD cytoperm) following the manufacturer’s instructions. These fixed cells were then stained using antibodies against several cytokines. As before, antibody cocktails used to stain the cells are given here - (TNF-α-FITC, IL-10-PE); (IFN-γ-FITC, IL-4-PE, IL-12-APC). The unstained, single colour stained and antibody cocktail stained cells were washed. Cells were maintained in permeabilization buffer and analyzed by flow cytometry. Compensation was performed if required using the single colour stained cells.

### Phagocytosis assay

Unstimulated and Fla^+Sia^/ds-Fla^+Sia^ matured moDCs were incubated for 30 min at 37°C with FITC-tagged latex beads (1:2000) as per the manufacturer’s protocols. Cells were washed three times with PBS to remove any surface-adherent beads and cells were analyzed by flow cytometry. Fluorescence at the FITC channel can be used to estimate the amount of internalized beads.

### Mixed lymphocyte reaction

PBMCs from donors were incubated in T75 flasks to remove the monocyte populations through plastic adherence as per Kizhakeyil et al (39). The monocyte-depleted cell population (responders) was stained with 5 µM CFSE dye followed by extensive washing. Responder cells were then incubated with ds-Fla^+Sia^/Fla^+Sia^ matured moDCs (stimulator) at 7:1 ratio (7X10^5^:1×10^5^) for 7 days in 24 well plates in RPMI medium supplemented with 10% human AB serum. After incubation, the cells were washed and then incubated with CD3-PE antibody for 30 min at 4°C to label the T cell population. These cells were washed with PBS and then analyzed by flow cytometry. T lymphocytes were detected based on CD3 positivity and CFSE-fluorescence within this population was measured. In a similar setup, the responder cells were again incubated with ds-Fla^+Sia^/Fla^+Sia^-matured-moDCs (stimulator) under identical conditions. After 7 days, all cells were washed and then stained with fluorescent tagged antibodies (CD3-PerCP, CD4-FITC and CD8-PE). Isotype controls, single-stained controls were included for detecting populations with appropriate compensation settings. The T lymphocytes populations were first detected through CD3 staining. Populations of T helper cells (CD3+CD4+) and cytotoxic T lymphocytes (CD3+CD8+) were separately detected. In another separate setup, T lymphocytes within the proliferating responder population was selected on the basis of CD3 positivity (CD3-V500) and any Treg cells produced within this population were detected through dual positivity upon staining with CD4-FITC and FoxP3-PE.

## Confocal Microscopy

### Binding of Fla^+Sia^ with TLR5, Siglec-9 in human monocyte derived DCs (moDCs)

Immature hmoDCs were incubated with FITC-Fla^+Sia^ at 4°C for 1 hour. Cells were washed and stained with anti-Siglec-9 antibody (1:1000) for further 1 hour at 4°C. Cells were washed, counterstained with PE-tagged secondary antibodies (1:2000) for 30 min at 4°C. These stained cells were fixed and then centrifuged on to coverslips before being mounted, sealed. The localization of Siglec-9 and FITC-Fla^+Sia^ on cell surfaces was analyzed by confocal microscopy. Colocalization resulted in overlap of the green (FITC-Fla^+Sia^) and red (Siglec-9) signals to give composite yellow colour in the merged images.

### Internalization of FITC-*Escherichia coli* by moDCs by confocal microscopy

FITC (0.1%) dissolved in 50 mM bicarbonate buffer, pH 8 was added to a suspension of PBS washed *Escherichia coli* ATCC 25922 and then incubated for 45 min at 37°C. The labelled bacteria were washed with PBS for 5 times to remove all traces of free FITC.

These FITC labelled *E. coli* ATCC 25922 were added at 10:1 ratio (or multiplicity of infection, MOI 10) to immature moDCs and flagellin matured moDCs and then incubated at 37°C for 30 min. Any unbound bacteria were removed by PBS washes. Fluorescence coming from extracellularly adherent bacteria was quenched by a brief staining with Trypan blue solution (0.02%). Cells were fixed with 2% paraformaldehyde and then transferred on to coverslips. Coverslips were mounted and observed through confocal microscopy to detect FITC labeled bacteria internalized by the cells. Images were acquired using Andor Spinning Disc Confocal microscope (Belfast,U.K.).

Acquired images were analyzed and the phagocytic index, which is the average of numbers of bacteria adhered/internalized per cell, was calculated.

### Genetic expression profiling by real time PCR

Total RNA from untreated or flagellin matured moDCs (1×10^6^) was extracted using RNeasy mini kit (Qiagen). Isolated RNA was quantified through its UV absorbance by a nanodrop and 2 μg was first treated with DNase I (Thermo Scientific) before being taken for the synthesis of cDNA (RevertAid First strand cDNA synthesis kit, Thermo Scientific). Primers for certain genes were obtained from Eurofins Genomics India Pvt. Ltd and given in **Table 1**. The expression status of these genes at the mRNA level was quantified through SYBR green based qRT-PCR using the DyNAmo Flash SYBR Green qPCR Kit (Invitrogen). The relative amounts of target mRNA in reference to 18S rRNA as internal control was estimated by Light Cycler 96 (Roche) software. Fold change in gene expression in ds-Fla^+Sia^ vs Fla^+Sia^ matured DCs were calculated using the comparative cycle threshold (CT) method.

**Table 1.**
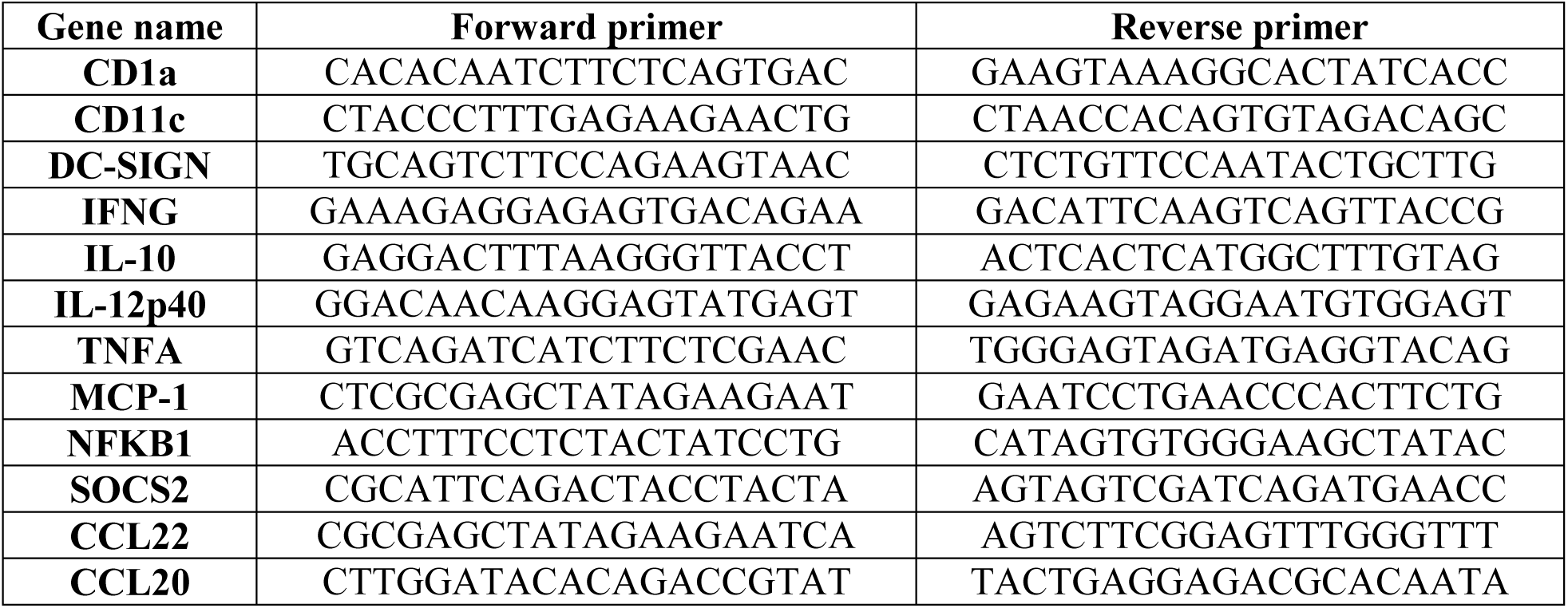
List of primers used in this study for qRT-PCR.

### Immunoblot and immunoprecipitation analysis

THP-1 derived DCs or moDCs (1 × 10^6^) stimulated with different doses of Fla^+Sia^/ds-Fla^+Sia^ for 24 hours were transferred into PBS buffer with protease, phosphatase inhibitors and lysed by sonication. Total protein content of cell lysates was quantified by BCA assay using a BSA standard. Equal amounts of lysates (70 μg), prestained protein ladder (Biorad) were resolved via SDS-PAGE. Proteins from gel were transferred to PVDF membranes using the wet transfer method.

In some cases, cells were resuspended in the cytosol isolation buffer with gentle vortexing (10 mM Tris-Cl, 10 mM NaCl, 1.5 mM MgCl_2_, 1 mM PMSF, 0.05% NP-40, pH 6.8) to allow partial lysis of cell membrane. The lysates were centrifuged at 1,000 g for 5 min and the supernatant containing the cytosolic extract was collected. The pellet was washed with chilled PBS, resuspended in nuclear isolation buffer (20 mM Tris-Cl, 137 mM NaCl, 1 mM CalCl_2_, 1 mM MgCl_2_, 1 mM PMSF, 1% NP-40, pH 8.0) and incubated for further 30 min along with intermittent vortexing. The samples were centrifuged and the supernatant containing the nuclear fraction was collected. As before, the protein content of cytosolic and nuclear fractions were quantified before subjecting to SDS-PAGE and western blotting.

Blots were blocked using BSA in TBS buffer and then were overnight incubated with primary antibody at 4°C with gentle shaking. Next day, blots were washed with TBS-Tween 0.1% buffer and appropriate secondary antibody dilutions in blocking buffer were added. Following 3 hours of room temperature incubation under gentle shaking, blots were again washed with TBS-Tween 0.1% buffer and binding of the secondary antibodies were probed using ECL substrate. Chemiluminescence signals were captured by Biorad ChemiDoc MP System or Azure c400 Visible Fluorescent Western Blot Imaging System.

The antibodies used in this study include - anti-TLR5 antibody (36–3900), anti-Siglec-9 antibody (703464), anti-MyD88 (MA516231) and anti-TRAF6 antibody (38–0900) from Invitrogen (Thermo Scientific); IRAK4 (4363S), p-p38 MAPK (#4511), p38 MAPK (#9212), p-Akt Ser473 (#4060), pan-Akt (#4691), p-ERK1/2 (#4377), ERK1/2 (#4695), p-JNK/SAPK (#4668), JNK/SAPK (#9252), β - Actin (#4970), Lamin B (#13435), NF-κB Pathway Sampler Kit (#9936), HRP-conjugated anti-rabbit secondary(#7047S) antibodies and anti-mouse secondary antibodies (#7076) from Cell signaling technology.

### NF-κB reporter assay

The activation and release of the transcription factor NF-κB p65 was assessed using a firefly luciferase-based NF-κB-reporter assay. The Luciferase NF-κB reporter plasmid was a gift from Dr. Anindita Ukil, University of Calcutta, Kolkata, India. All THP-1 derived DCs were transfected with the NF-κB reporter while some were co-transfected with the reporter and Siglec-9/TLR5 siRNA and some with only Lipofectamine Plus. After 24 hours of transfection, some cells containing only the reporter were further treated with inhibitors (TH1020/NSC-87877). These transfected and inhibitor treated cells were then stimulated with Fla^+Sia^ in a 96 well plates for 24 hours. Cells were lysed and then processed for luciferase assay at 48 h post-transfection as per the Luciferase assay kit. Readings were taken in a plate multireader.

### ELISA based determination of secreted cytokines

THP1 derived DCs (1.5X10^6^) were stimulated with Fla^+Sia^ following TLR5/Siglec-9 knockdown or inhibitors treatment (TH1020/NSC-87877). Culture supernatant from these cells was collected and the levels of secretory cytokines-IFN-γ (E-EL-H0108) and IL-12 (E-EL-H0150) were determined by ELISA kits following manufacturer’s instructions (Elabscience, Texas, USA).

## Statistical Analysis

Data presented here are mean values calculated from at least three independent experiments. For two groups of samples, two-tailed Student’s t-test was used to derive the statistical significance of the recorded data. One-way analysis of variance (ANOVA) was used when more than two groups were involved along with pair wise multiple comparison procedures via Tukey test where p < 0.05 was deemed to be statistically significant. Error-bars in the figures represent mean ± standard error of the mean (SEM) values derived from three independent experiments. Significant differences were defined as ns=p > 0.05, *=p ≤ 0.05, **=p ≤ 0.01, ***=p ≤ 0.001, and ****=p ≤0.0001. Data were calculated and analyzed using GraphPad Prism version 6.01.

## Results

### The purification and characterization of PA flagellin

It has been previously established that a sialylated form of *Pseudomonas aeruginosa* (PA) can be obtained when the bacteria is cultured in a medium containing a sialic acid source like fetal calf serum (FCS) (4–6, 34). Previously, through affinity chromatography, two sialic acid-specific lectins were used to isolate and identify a few sialic acid containing proteins from these sialylated bacteria (PA^+Sia^) (6). An important sialylated protein identified in this manner was the PA flagellin (6).

Based on this background, a lectin-based pull down assay was performed to confirm once more that sialic acid-specific lectins can bind with PA flagellin. The PA14 strain were cultured overnight in 10% FCS supplemented Tryptic Soy Broth (TSB) medium, following which PA^+Sia^ were pelleted, washed and then lysed by gentle sonication. This bacterial lysate was incubated with a sialic acid-specific lectin – *Sambucus nigra* agglutinin (SNA) conjugated to agarose beads. Those proteins which remained bound to the SNA-agarose beads were dissolved into SDS containing sample buffer and then resolved via SDS-PAGE. The gel was stained with Coomassie Blue and destained to reveal several fine bands along with two distinct bands near 34 kDa and one prominent band near 47 kDa (**Figure 1A**). Bovine submaxillary mucin (BSM), which was used as a positive control, showed similar double bands near 34 kDa. This suggests that these distinct protein bands come from the SNA lectin itself. The distinct band near 47 kDa was excised, reduced, alkylated and then digested with trypsin before being subjected to MALDI TOF MS/MS. Through peptide mass fingerprinting, this band was identified to be PA flagellin type B (**Figure 1A**). This corroborates with the previously reported data that sialic acid specific lectin SNA can recognize and bind to flagellins isolated from PA^+Sia^.

**FIGURE 1.**
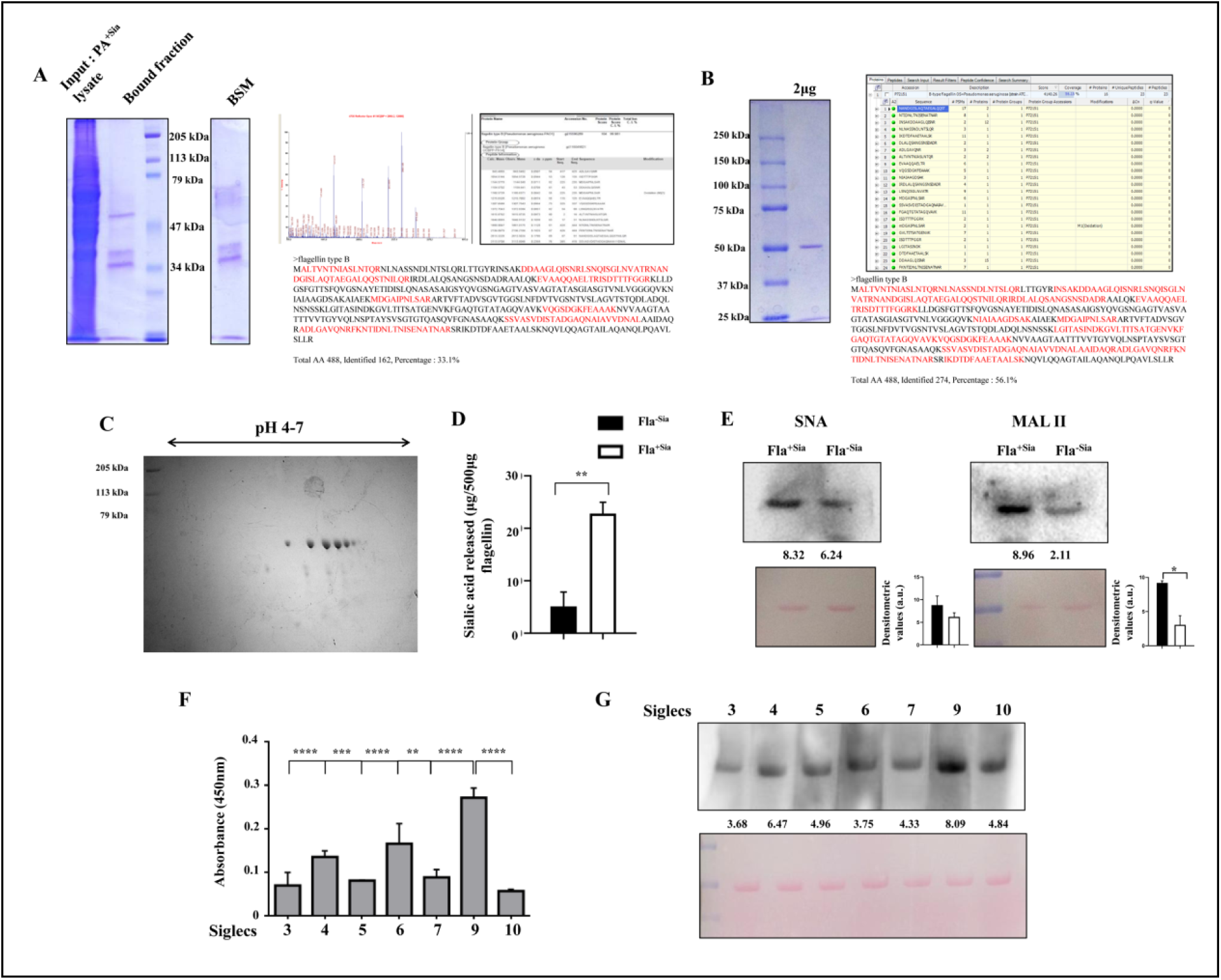
Evaluation of the sialylation status of flagellin isolated from PA^+Sia^ **(A)** The sialic acid specific – *Sambucus nigra* (SNA) lectin conjugated to agarose beads was used to isolate sialoglycoproteins from PA^+Sia^ lysate (100 µg) by a lectin-based pulldown assay. Highly sialylated bovine salivary mucin (BSM) was used as a positive control. In the case of PA^+Sia^ lysate, the lectin bound sialylated proteins were resolved in 10% SDS-PAGE and visualized by Coomassie Blue staining. Staining revealed several fine bands along with a prominent band near 47 kDa and two bands near 34, 37 kDa. The presence of similar bands in the BSM control indicated that these 34, 37 kDa bands come from the SNA-agarose beads. The band near 47 kDa was excised and processed for *in gel* trypsin digestion. The protein band was reduced, alkylated then digested by trypsin and subjected to MALDI TOF MS/MS. The generated spectra were searched against a eubacterial protein database by the MASCOT search engine utilizing the Basic Local Alignment Tool (BLAST) of ABI GPS Explorer software, version 3.6. Through peptide mass fingerprinting, the protein band was identified as flagellin. The MS/MS spectra with each tryptic fragments denoted by respective m/z values and the BLAST search results are shown. The peptide fragments within the flagellin sequence which were detected via MALDI TOF MS/MS have been marked with red colour. **(B)** The isolated, purified Fla^+Sia^ protein (2 μg) resolved to a single band in a 10% SDS-PAGE gel. This single band was carefully excised and subjected to *in gel* trypsin digestion as before. The peptide digest was analyzed via ESI MS/MS mode by the LTQ Orbitrap mass spectrometer. The generated spectra was searched against the eubacterial protein database compiled from Uniprot using the Sequest HT Proteome Discoverer 1.4 software (Thermo Fisher Scientific) utilizing the MASCOT search engine. All peptide fragments among the flagellin sequence which have been identified in the tryptic digest have been denoted with red colour. **(C)** Purified Fla^+Sia^ (25 μg) was passively rehydrated into pH 4–7 IPG strips and resolved via isoelectric focussing. The resolved strips were placed over the resolving gel in a10% SDS-PAGE gel and fixed using melted agarose. The protein sample was resolved in the second dimension through electrophoresis. The 2D gels were stained using Coomassie Blue to reveal that the single band protein had split into seven isoforms**. (D)** The sialic acid content of Fla^+Sia^/Fla^-Sia^ (500 µg) were estimated and compared using the fluorimetric acetyl acetone method following Shukla et al (1982). Following the acetyl-acetone reaction, sialic acids released from flagellin proteins formed a fluorigenic product which were quantified via a fluorimeter (Ex-410nm, Em-510nm). The amount of sialic acids present in each sample was calculated by comparing with a prepared standard dilution of synthetic N-acetyl Neuraminic acid. Data from three independent experiments were used to prepare the bar diagram shown here. Significance is represented by *p ≤ 0.05, **p ≤ 0.01, ***p ≤ 0.001, and ****p ≤ 0.0001. **(E)** Fla^+Sia^/Fla^-Sia^ (5μg) resolved via SDS-PAGE was transferred onto PVDF blots via wet transfer method. These blots were probed with biotinylated sialic acid binding lectins – *Sambucus nigra* agglutinin (SNA) and *Maackia amurensis* agglutinin (MAA II). The blots were washed and incubated with secondary avidin-HRP and the extent of lectin binding was determined by developing the blot using chemiluminescent substrates. The Ponceau S stained blots are provided to account for equal protein loading. Densitometric data from three independent experiments were used to prepare the bar diagram shown here. Representative blots from one experiment are shown here. Significance is represented by *p ≤ 0.05, **p ≤ 0.01, ***p ≤ 0.001, and ****p ≤ 0.0001. **(F)** Fla^+Sia^ was used to coat a 96 well ELISA plate. A panel of Siglec-Fc chimera recombinant proteins (Siglec-3,4,5,6,7,9 and 10-Fc) was diluted and incubated in these Fla^+Sia^ coated wells. The wells were then washed and incubated with anti-human IgG Fc-specific peroxidase secondary antibodies. The extent of Siglec-Fc remaining bound to the Fla^+Sia^ coated wells were detected using TMB substrate. The colour which was generated from the bound secondary antibodies was measured as absorbance at 450nm in a plate reader. Data from three independent experiments were used to prepare the bar diagram shown here. Significance is represented by *p ≤ 0.05, **p ≤ 0.01, ***p ≤ 0.001, and ****p ≤ 0.0001. **(G)** As before, Fla^+Sia^/Fla^-Sia^ (5μg) were resolved in SDS-PAGE, transferred to blots and probed using the same Siglec-Fc panel. The blots were washed and then incubated with secondary anti-human IgG Fc region specific peroxidase antibodies. The extent of Siglec binding was directly visualized by developing the blots in a chemidoc using a chemiluminescent substrate. As before, Ponceau S stained blots are supplied to account for equal loading. One representative blot from three independent experiments performed is shown here.

The PA14 bacteria were again cultured in a large volume of FCS supplemented Tryptic Soy Broth (TSB) to obtain a high yield of PA^+Sia^. Flagellin was isolated from these PA^+Sia^ following the method described by Campodónico et al (37) which involved gentle shearing of extruded surface proteins of the bacteria followed by salting out. After dialysis, the purified flagellin proteins were incubated with agarose-bound polymyxin B sulfate beads to remove any contaminating endotoxins and the endotoxin content was confirmed to be below 0.1 EU/mL. The isolated protein was resolved into a single band near 47 kDa marker in SDS-PAGE, indicating its purity **(Figure 1B and Supplementary Figure S1)**. The molecular mass of the isolated protein was also similar to the theoretical molecular weight of PA flagellin type B (49 kDa) (10).This isolated, single-band protein was processed through *in gel* trypsin digestion and the digest was analyzed via ESI-MS/MS by LTQ-Orbitrap. The isolated protein was confirmed to be PA flagellin **(Figure 1B and Supplementary Fig S2)**. Thus, the process of isolating flagellin proteins from PA^+Sia^ in a pure, native state was standardized.

Flagellins purified from PA^+Sia^ were resolved by isoelectric focusing along a pH 4-7 gradient and then resolved in the 2^nd^ dimension. The 2D gel was stained by Coomassie Blue which revealed that the protein which resolved into a single band in SDS-PAGE, had resolved into seven distinct spots (**Figure 1C**). This indicates the presence of microheterogeneity, which is a possible indicator that flagellin exists in several altered glycosylation states **(Figure 1C)**. These seven distinct spots were carefully excised and again identified by MALDI-TOF based peptide mass fingerprinting. All seven spots were confirmed to be PA flagellin through MALDI TOF MS/MS (**Supplementary Fig S3 and S4)**.

### Establishment of the sialylation status of PA flagellin

Flagellin was separately isolated from sialylated (PA^+Sia^) and non-sialylated PA (PA^-Sia^). Using the fluorimetric acetylacetone assay method (38), the sialic acid content of equal amounts (500 µg) of isolated flagellin proteins from PA^+Sia^ and PA^-Sia^ were quantified and compared. The sialic acid content of flagellin from PA^+Sia^ (Fla^+Sia^) was estimated to be significantly greater by nearly 4.5 times (22.82 ± 1.24 μg) than the flagellin from PA^-Sia^ (Fla^-Sia^) (5.100 ± 1.58 μg) **(Figure 1D)**. The presence of sialic acids in Fla^+Sia^ was also checked by lectin binding. Equal amounts of Fla^+Sia^ (5μg) were resolved via SDS-PAGE and then transferred to polyvinylidene difluoride (PVDF) membranes. These blots were incubated with biotinylated lectins which specifically bind to sialic acids-*Sambucus nigra* agglutinin (SNA) and *Maackia amurensis* agglutinin type II (MAA II). These blots were washed and then incubated with avidin-HRP. Both lectins were observed to preferentially bind with Fla^+Sia^ compared to Fla^-Sia^ **(Figure 1E)**. Interestingly, the lectin MAA II showed nearly 4 fold higher binding with Fla^+Sia^ than Fla^-Sia^. Since, MAA II preferentially binds with α2-3 linked sialic acids, this indicates that Fla^+Sia^ contains a greater presence of sialic acids with such linkages.

Siglecs are Immunoglobulin-type (I-type) lectins commonly expressed by immune cells which specifically recognize and bind to sialic acids (17, 18). Therefore, a panel of commercially available Siglec recombinant proteins conjugated to human Fc regions was selected in order to check for any possible interactions with Fla^+Sia^. This panel covers most Siglecs normally expressed in human immune cells (30). ELISA plates were coated with Fla^+Sia^ and then incubated with these Siglec proteins. The wells were washed to remove any unbound proteins and later incubated with anti-human-Fc peroxidase conjugated secondary antibodies. After washing these wells, TMB substrate was added which reacted with bound peroxidase-conjugated secondary antibodies to give a color product. Thus, the extent of Siglec binding to Fla^+Sia^ could be detected by checking the absorbance at 450 nm. Here, the Siglecs-4, 6, and 9 were found to bind to the Fla^+Sia^ coated wells **(Figure 1F).** Out of all the tested Siglecs, Siglec-9 was found to bind most strongly with Fla^+Sia^.

As before, Fla^+Sia^/Fla^-Sia^ blots were incubated with the same panel of Siglec-Fc recombinant proteins. The blots were washed and then further incubated with anti-human-Fc-peroxidase secondary antibodies. Finally, the blots were washed and visualized in a chemidoc after the addition of a chemiluminescent substrate. As before, Siglec-9 was found to strongly bind with Fla^+Sia^, while Siglecs-4 and 10 showed some binding as well **(Figure 1G)**.

### The removal of sialic acids from flagellin

In order to further confirm the presence of sialic acids in PA flagellin, the isolated Fla^+Sia^ were treated with *Arthrobacter ureafaciens* neuraminidase which should remove any sialic acid residues present in the protein. This sialidase treated, desialylated Fla^+Sia^ is denoted as ds-Fla^+Sia^.

Equal amounts (20 µg) of sialidase treated or untreated flagellin was resolved in IPG strips via isoelectric focusing in a 3-10 pH gradient. After staining and destaining, the resolution profiles were compared. As observed before with 2D electrophoresis (**Figure 1C**), Fla^+Sia^ resolved into multiple bands in the pH gradient (**Figure 2A**). Interestingly, the sialidase treated flagellins show a slightly altered resolution profile compared to the untreated flagellin (ds-Fla^+Sia^ vs Fla^+Sia^) (**Figure 2A**). Sialic acids are negatively charged acidic sugars which contribute to the isoelectric point (pI) of proteins. Thus, a removal of sialic acids will result in a change in the protein pI, which affects the protein resolution in isoelectric focusing. Here, the ds-Fla^+Sia^ bands travelled further towards the basic pole (pH 10) compared to Fla^+Sia^. However, the observed shift in pI was not drastic, which again supports the existing knowledge that Fla^+Sia^ is not a highly glycosylated protein (10).

**FIGURE 2.**
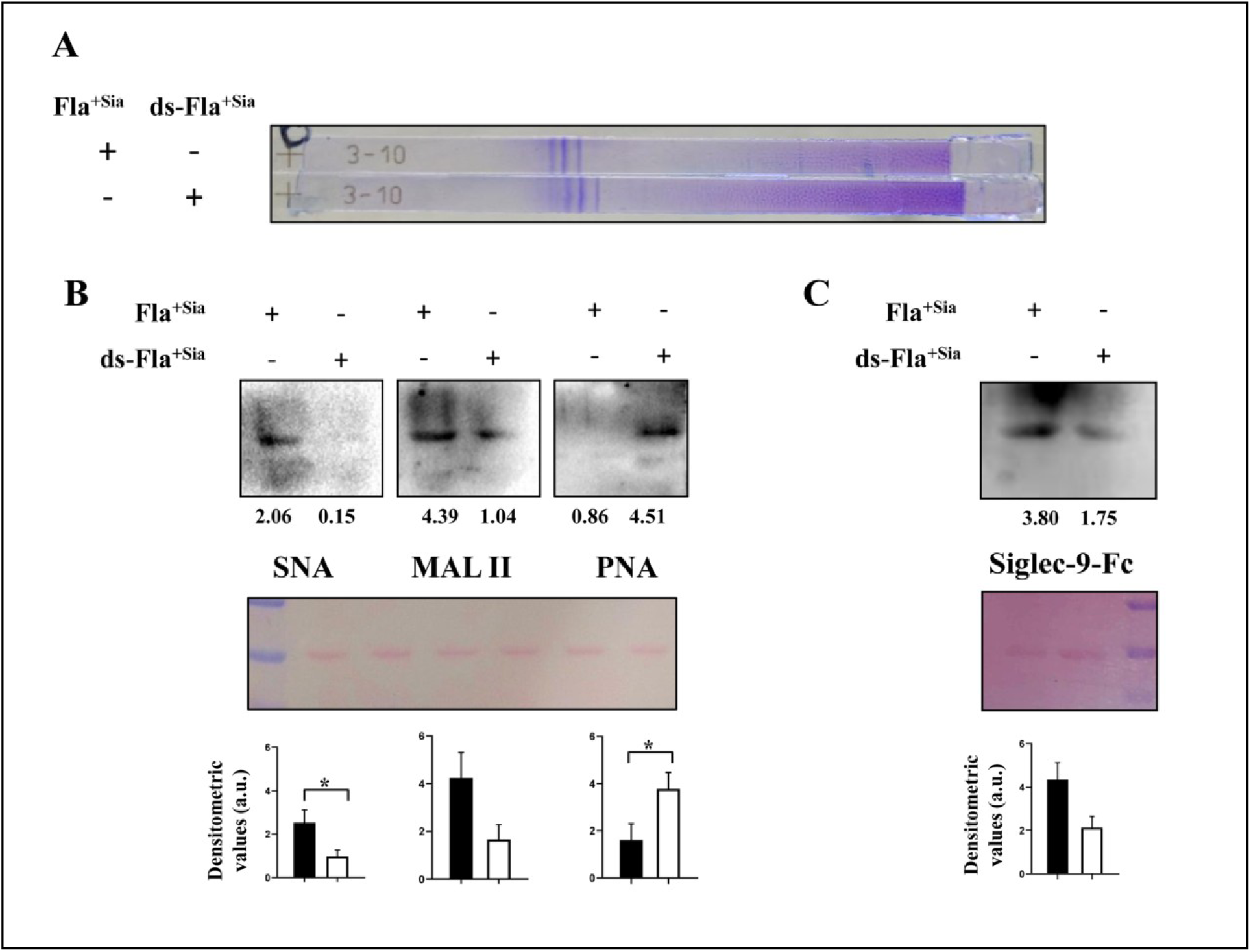
The confirmation of flagellin sialylation status through neuraminidase treatment. **(A)** Purified Fla^+Sia^ or desialylated Fla^+Sia^ (ds-Fla^+Sia^) (20 μg) were passively rehydrated into pH 3-10 IPG strips. The proteins were resolved by isoelectric focusing following the manufacturer’s instructions. These strips were stained by Coomassie blue and then destained before being dried. IPG strips were observed to detect any changes in protein resolution upon sialidase treatment. Representative strips selected from the two independent experiments performed are shown here. **(B)** As before, Fla^+Sia^/ds-Fla^+Sia^ (5μg) were resolved in SDS-PAGE, transferred on to PVDF blots and probed with biotinylated sialic acid binding lectins – SNA, MAA II ((5 μg/mL). The blots were washed and then incubated with avidin-HRP. The extent of lectin binding was determined by developing the blots in a chemidoc using a chemiluminescent substrate. Additionally, the galactose specific lectin – peanut agglutinin (PNA) was also used (10 μg/mL). Upon sialidase treatment, the removal of sialic acid residues at the glycan termini exposes the underlying galactose residues. PNA recognizes and binds these exposed galactose residues indicating the successful removal of sialic acids. Densitometric data from three independent experiments were used to calculate the mean band intensities which are shown in these graphs representing the binding studies. One representative blot per experiment is shown here. Significance represented by *p ≤ 0.05, **p ≤ 0.01, ***p ≤ 0.001, and ****p ≤ 0.0001. **(C)** In a same set up, Fla^+Sia^/ds-Fla^+Sia^ blots were probed with Siglec-9-Fc chimera proteins. The blots were washed and then incubated with anti-human IgG Fc specific antibodies. Any change in Siglec-9 interaction with flagellin was detected by developing the blot in a chemidoc. As previously described, Ponceau S stained blots are shown to account for equal protein loading across lanes. Densitometric data from three independent experiments were used to calculate the mean band intensities which are shown in these graphs representing the binding studies. One representative blot per experiment is shown here. Significance represented by *p ≤ 0.05, **p ≤ 0.01, ***p ≤ 0.001, and ****p ≤ 0.0001.

Equal amounts of Fla^+Sia^ and ds-Fla^+Sia^ (5 µg) were resolved via SDS-PAGE and transferred to PVDF blots. These blots were then incubated with sialic acid-specific lectins and Siglecs to check if desialylation had caused any change in their binding. The sialidase treatment of Fla^+Sia^ abrogated the binding of both SNA and MAA II **(Figure 2B)**. Enzymatic desialylation removes the terminal sialic acid residues in the glycan chains and exposes the underlying galactose residues; which can be recognized by the peanut agglutinin (PNA) lectin. As expected, PNA lectin showed a strong binding with only the sialidase-treated flagellin (ds-Fla^+Sia^), confirming that successful desialylation has taken place **(Figure 2B)**. The sialidase treatment of Fla^+Sia^ also reduced the binding of Siglec-9, indicating the removal of sialic acid residues which are the ligands for Siglec-9 **(Figure 2C)**. Together, these observations further establish the presence of sialic acids in Fla^+Sia^ as well as confirm its removal.

### Monocyte-derived dendritic cells (moDCs) bind to sialylated flagellin through TLR5 and Siglec-9

As a highly conserved Pathogen Associated Molecular Pattern (PAMP), flagellins are detected by the Toll Like Receptor-5 (TLR5) on the cell surface, which leads to the generation of strong pro-inflammatory responses (40, 41). Thus, due to their strong antigenicity, flagellin stimulus can be used to induce dendritic cell (DC) maturation (25, 42–44). Here, we wanted to assess if the sialic acids present in flagellin have any impact on this critical process. Towards this purpose, monocytes were isolated from human PBMCs and stimulated to differentiate into immature monocyte derived DCs (moDCs). The cells were induced to differentiate through stimulation with rhGM-CSF (recombinant human granulocyte-macrophage colony stimulating factor) and rhIL-4 (recombinant human interleukin-4) as per standard protocols (35). The generation of immature moDCs was monitored through flow cytometry by checking for changes in the expression of cell markers, like the reduction in the monocyte marker –CD14 and the increase in markers like CD80, CD86, DC-SIGN (CD209) etc (see **Supplementary Figure S5**) (35). These immature moDCs were incubated with anti Siglec-5 and anti Siglec-9 antibodies and counterstained with the fluorophore phycoerythrin (PE) conjugated secondary antibodies. The expression status of Siglecs-5 and 9 on the surface of immature moDCs was checked by flow cytometry. Compared to Siglec-5, a robust expression of Siglec-9 was detected in the cells (**Figure 3A**).

**FIGURE 3.**
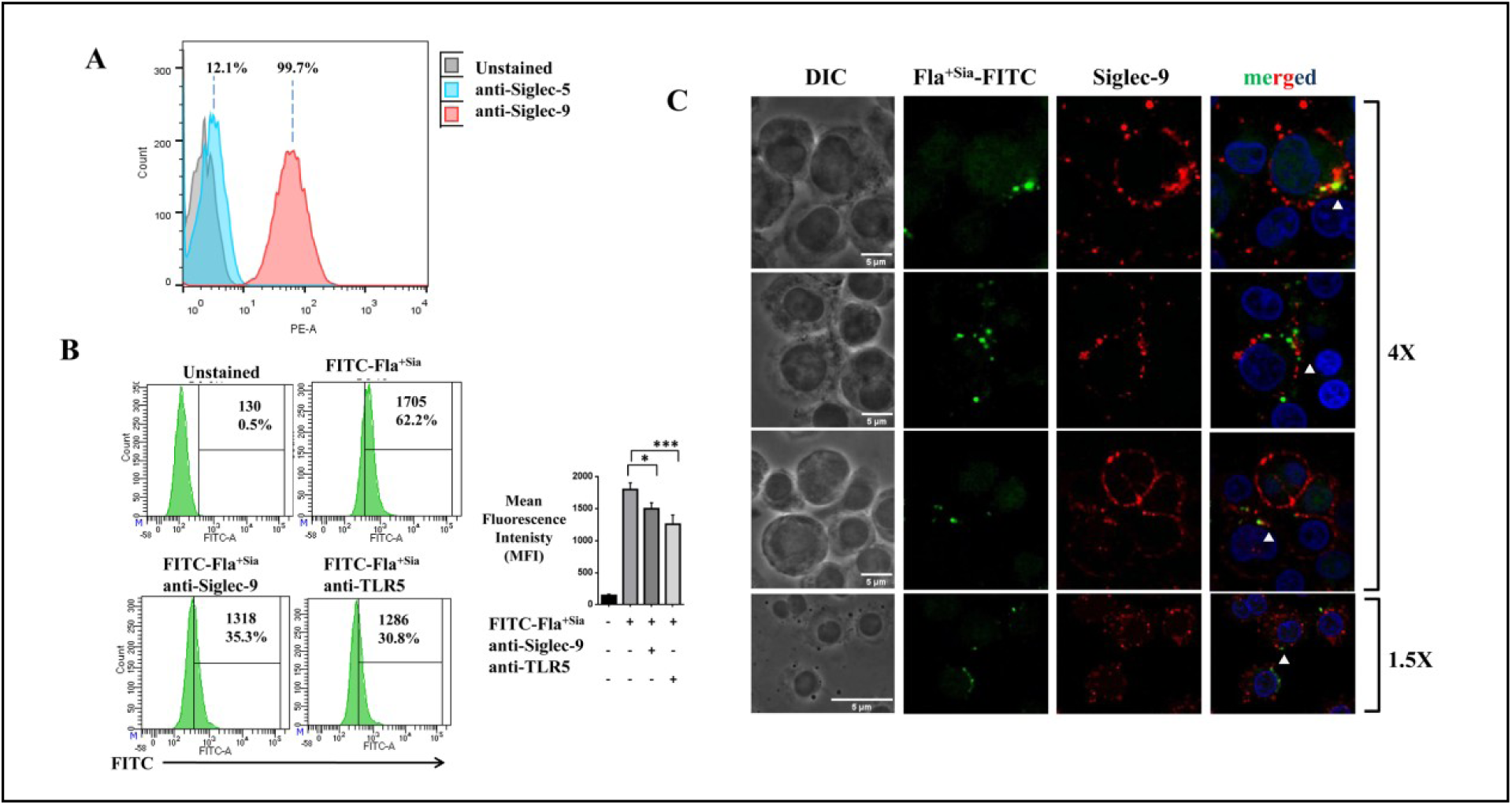
Fla+Siainteracts and binds with TLR5 and Siglec-9 on the dendritic cell surface. **(A)** Human monocyte derived dendritic cells (hmoDCs) were generated and these immature cells were stained with anti Siglec-5 and anti Siglec-9 antibodies (1:1000) respectively for 1 hour at 4°C. Cells extensively washed with PBS and then counterstained with phycoerythrin (PE) labelled secondary antibodies. Cells were again washed and finally analyzed by flow cytometry to detect the Siglecs expression on the cell surface. The experiment was repeated independently twice and a representative histogram is shown here. **(B)** The binding of FITC tagged Fla^+Sia^ by moDCs was evaluated by flow cytometry. Immature moDCs were pre-treated with anti Siglec-9 or anti TLR5 antibodies (1:1000). Untreated and antibody-blocked moDCs (1×10^6^) were incubated for 1 hour with FITC-Fla^+Sia^ (2μg) to allow for binding at 4°C. These cells were washed with PBS and then analyzed by flow cytometry. The extent of Fla^+Sia^ binding was measured from the FITC signal associated with untreated/treated DCs. Data from three independent experiments is used to prepare the bar diagram and representative histograms from one such experiment is shown here. Significance is represented by *p ≤ 0.05, **p ≤ 0.01, ***p ≤ 0.001, and ****p ≤ 0.0001. **(C)** Immature moDCs were incubated with FITC-Fla^+Sia^ at 4°C for 1 hour followed by PBS wash. These cells were then stained with anti-Siglec-9 antibodies (1:1000) for another 1 hour at 4°C. Cells were counterstained with PE-tagged secondary antibodies (1:2000) followed by fixation before being mounted and sealed. The localization of PE stained Siglec-9 and FITC-Fla^+Sia^ on cell surfaces were visualized by confocal microscopy and regions of colocalization (red-green) showed up as yellow in the merged images. Images taken at two different magnifications (4X and 1.5X) are shown here. The regions of colocalization are indicated with white arrows. Representative images selected from three independent experiments are shown here.

The previous experiments (**Figure 1G, 1F**) have established that Siglec-9 strongly binds with Fla^+Sia^. Additionally, immature human moDCs have been reported to express the flagellin cognate receptor - TLR5 (45). Therefore, we next checked if Fla^+Sia^ interact with TLR5 as well as Siglec-9 present on the immature moDC surfaces. Towards this purpose, Fla^+Sia^ were first labeled with fluorescein isothiocyanate (FITC) by incubating with FITC dissolved in bicarbonate buffer. Following incubation, any unbound FITC was removed from the protein solution through extensive buffer exchange in a centrifugal filter. Generated immature moDCs were pre-treated with anti Siglec-9 and anti TLR5 antibodies respectively. These antibody-blocked and untreated DCs were incubated with FITC-Fla^+Sia^ to check for binding. Flow cytometry revealed that the extent of FITC-Fla^+Sia^ binding with moDCs (MFI = 1796 ± 53.35) (**Figure 3B)** was reduced when cells were pretreated with blocking antibodies against TLR5 (MFI = 1257 ± 84.30). Interestingly, Fla^+Sia^ binding was also reduced when cell surface Siglec-9 was blocked by antibody treatment (MFI = 1596 ± 116.4). This indicates that Fla^+Sia^ interact with both TLR5 and Siglec-9 on the moDC surface.

The association between FITC-Fla^+Sia^ and cell surface Siglec-9 was next visualized by confocal microscopy. Non-permeabilized moDCs incubated with FITC-Fla^+Sia^ were stained with anti-Siglec-9 antibodies followed by PE tagged secondary antibodies. Confocal microscopy revealed a few areas of colocalization between FITC-Fla^+Sia^ and PE-stained Siglec-9 on cell surfaces (**Figure 3C**). The areas of colocalization could be detected through the overlap of red and green fluorescence, giving a yellow signal in the composite images. Taken together, these experiments show that Fla^+Sia^ can interact and bind with both TLR5 as well as Siglec-9 on the moDC cell surface.

### The impact of Fla^+Sia^ stimulation on moDC maturation

Generally, the TLR5-based detection of flagellin leads to MAP Kinase signaling, NF-κB activation, pro-inflammatory cytokines secretion and the upregulation of MHC and costimulatory molecules (40). Interestingly, Fla^+Sia^ was observed to interact with both pro-inflammmatory TLR5 as well as the immune-inhibitory Siglec-9 on the immature moDC surface. Thus, it was crucial to check if these interactions (particularly Sigle-9) had any effect on the DC maturation. Accordingly, immature moDCs were incubated with increasing doses of sialylated or desialylated flagellins (Fla^+Sia^/ds-Fla^+Sia^). After 24 hours of incubation, the expression of costimulatory/maturation markers (CD86, CCR7, HLA-DR) and DC-specific markers (CD83, CD11c) were estimated to determine the DC maturation status.

The immature moDCs stimulated with increasing concentrations of ds-Fla^+Sia^ showed an increased expression of markers like CD83, CD86, HLA-DR (**Figure 4A).** Interestingly, stimulation with same doses of Fla^+Sia^ resulted in comparatively reduced expression of these same markers. At the highest concentration of treatment (5µg), moDCs treated with of Fla^+Sia^ show a significant reduction in the expression levels of the maturation markers-CD86, CCR7, HLA-DR and DC-specific markers-CD83, CD11c as compared to ds-Fla^+Sia^ treated ones (**Figure 4A**). At 5 µg stimulated ds-Fla^+Sia^ versus Fla^+Sia^ stimulated DCs, the MFI values of DC-specific marker CD83 was (507.7 ± 30.89 vs 356.0 ± 11.27); major histocompatibility complex (MHC) II cell surface receptor (HLA-DR) was (1094 ± 91.51 vs 735.7 ± 22.28); CD86 – a costimulatory receptor necessary for T cell activation was (553.7 ± 18.77 vs 366.3 ± 29.74); and so on.

**FIGURE 4.**
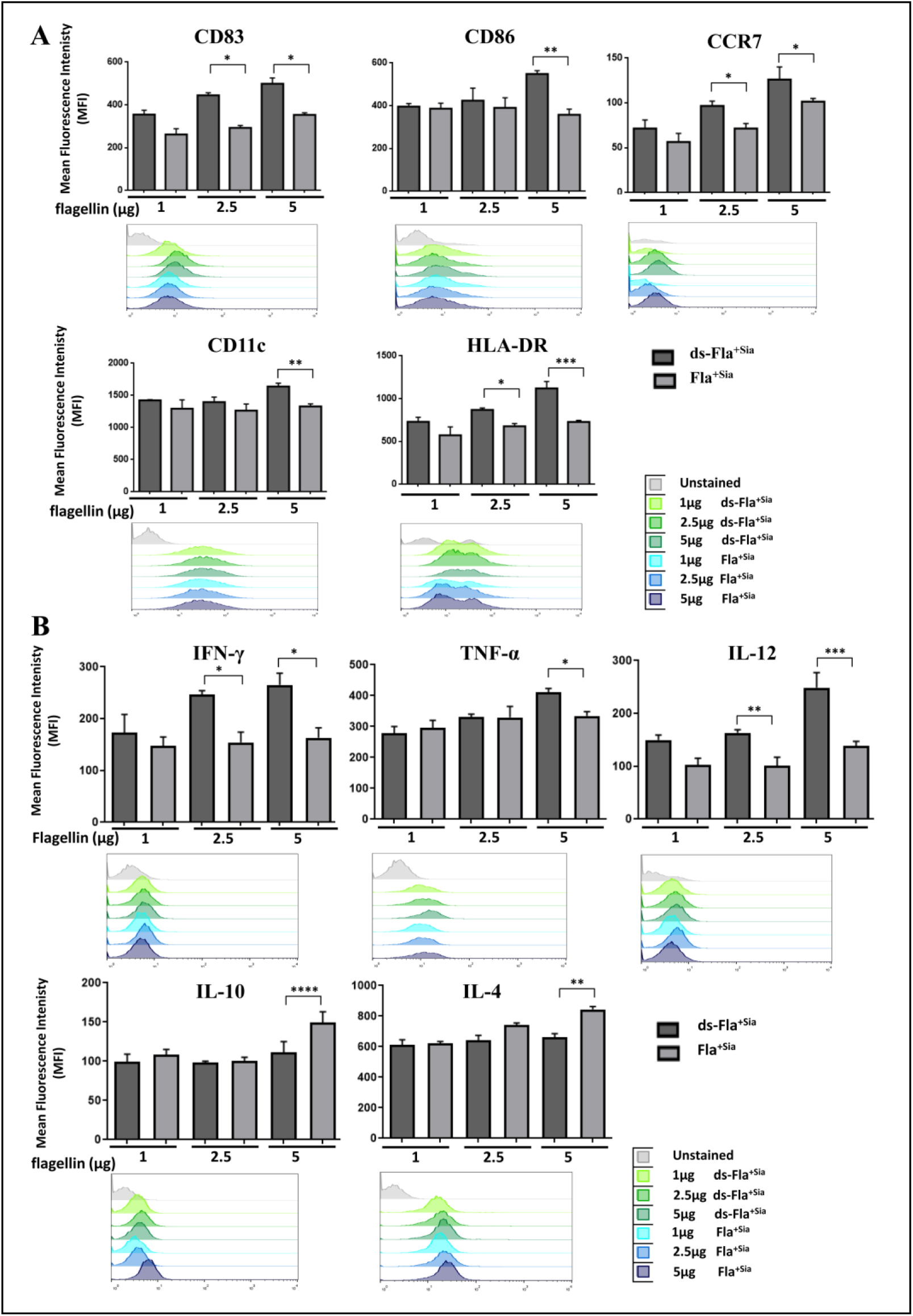
Differential maturation profiles of Fla^+Sia^/ds-Fla^+Sia^ stimulated dendritic cells. **(A)** The expression of maturation markers and co-stimulatory receptors on moDCs matured with different doses of Fla^+Sia^/ds-Fla^+Sia^ (1 μg, 2.5 μg, 5 μg) for 24 hours was estimated by flow cytometry. The treated cells were stained with the cocktails of appropriate fluorophore-conjugated antibodies for 30 min at 4°C followed by PBS wash. Unstained, single-color stained samples and cocktail stained cells were then analyzed in a flow cytometer. Appropriate compensation settings were used when required. Data from at least three independent experiments were used to prepare bar diagrams showing the changes in the surface marker expression. Representative histograms showing shifts in the stained populations are shown. Significance is represented by *p ≤ 0.05, **p ≤ 0.01, ***p ≤ 0.001, and ****p ≤ 0.0001. **(B)** Similarly treated matured DCs were fixed and permeabilized following the manufacturer’s instructions. These cells were then stained using cocktails of fluorophore labeled antibodies against pro-inflammatory (IFN-γ,TNF-α,IL-12) and anti-inflammatory cytokines (IL-4, IL-10) cytokines. Cells were then washed and maintained in permeabilization buffer. Unstained, single-color stained samples and stained cells were analyzed by flow cytometry to detect the intracellular levels of cytokines. Adequate compensation settings were used in the case of multicolour staining. Bar diagrams prepared from more than three independent experiments depict the change in observed in cytokine expression. Representative histograms showing the shifts in stained populations are shown. Significance is represented by *p ≤ 0.05, **p ≤ 0.01, ***p ≤ 0.001, and ****p ≤ 0.0001.

The intracellular levels of cytokines produced in Fla^+Sia^/ds-Fla^+Sia^ stimulated moDCs were also determined **(Figure 4B**). Matured cells responded with increased levels of pro-inflammatory cytokines, particularly, IFN-γ, IL-12 across the varying concentration of ds-Fla^+Sia^ stimuli compared to Fla^+Sia^. In the case of TNF-α, the difference was pronounced at the highest dose of flagellin (5 μg). At 5 μg stimulation, the MFI values denoting the pro-inflammatory cytokines secretion by ds-Fla^+Sia^ compared to Fla^+Sia^ were – IFN-γ (MFI: 265.3 ± 38.89 vs. 163.0 ± 33.45); IL-12 (249.0 ± 28.51 vs. 142.0 ± 11.36) and TNF-α (413.7 ± 14.47 vs. 339.3 ± 15.89). In contrast, at the highest dose of stimulation (5 μg), the Fla^+Sia^ maturated moDCs exhibited an increased production of anti-inflammatory cytokines like IL-10 (MFI: 149.7 ± 13.50 vs. 111.0 ± 27.53) and the pleiotropic sytokine IL-4 (MFI: 843.3 ± 34.02 vs. 660.3 ± 43.55) as compared to ds-Fla^+Sia^-stimulated moDCs (**Figure 4.2B**). Thus, moDCs stimulated with Fla^+Sia^ exhibit an interesting deviation from the predominantly pro-inflammatory response of ds-Fla^+Sia^ matured moDCs. Taken together, these observations indicate that the DCs matured with Fla^+Sia^ exhibit a different maturation profile compared to ds-Fla^+Sia^ stimulation.

### The effectivity of DCs stimulated with Fla^+Sia^

Immature DCs are also functionally different from mature DCs. Immature DCs exhibit profuse dendritic processes along with high phagocytic activity. This phagocytic activity is reduced after maturation (20).

Therefore, we next compared the phagocytic ability of immature, Fla^+Sia^ or ds-Fla^+Sia^ matured DCs by checking their ability to internalize FITC-tagged latex beads. Immature moDCs phagocytosed the highest amount of FITC-beads leading to high MFI values (MFI = 1965 ± 49.57) (**Figure 5A**). Fla^+Sia^ matured DCs were also found to retain their phagocytic activity to a greater extent (1701 ± 89.25), while it was diminished in the case of ds-Fla^+Sia^ matured DCs (1017 ± 119.3).

**Figure 5.**
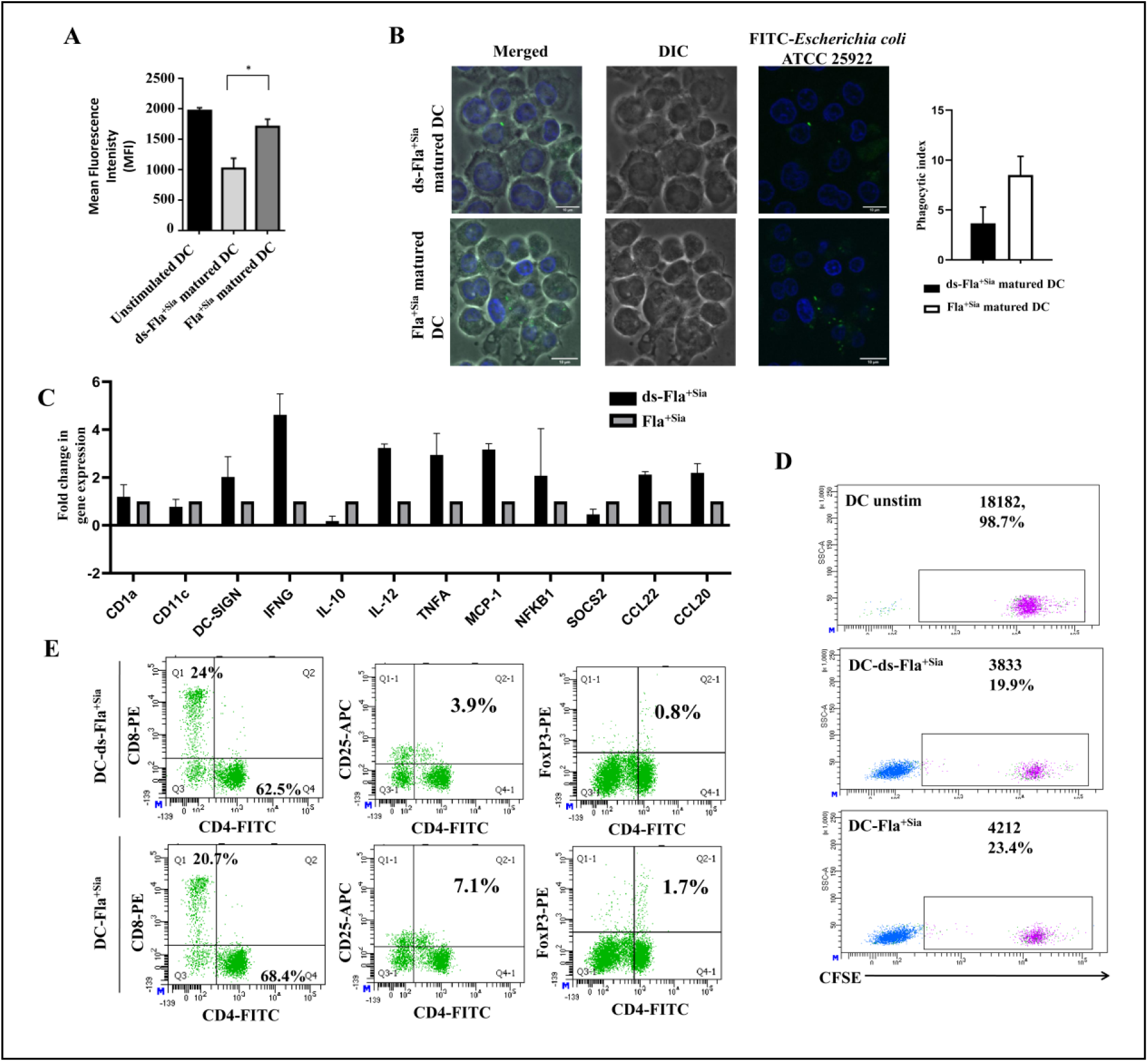
Functional activities and T cell proliferation ability of flagellin matured DCs. **(A)** The phagocytic property of immature moDCs and Fla^+Sia^/ds-Fla^+Sia^ matured moDCs was compared by the uptake of FITC-latex beads. The moDCs were incubated for 30 min at 37°C with FITC-tagged latex beads (1:2000) followed by extensive PBS washes to remove all unbound beads. These cells were then analyzed through flow cytometry to detect the intracellular FITC signal. Data from at least three independent experiments are represented in the bar diagram. Significance is represented by *p ≤ 0.05, **p ≤ 0.01, ***p ≤ 0.001, and ****p ≤ 0.0001. **(B)** The Fla^+Sia^/ds-Fla^+Sia^ matured moDCs were incubated with FITC labeled *Escherichia coli* ATCC 25922 at 1:10 ratio at 37°C for 30 min. All unbound bacteria were removed through extensive PBS wash, the cells were fixed and then visualized by confocal microscopy. Bacteria associated with DCs were observed as green dots (FITC signal). The internalized bacteria were counted to estimate the phagocytic index (number of bacteria/cell). Data from at least 40 cells were used to prepare the bar diagram. Significance is represented by *p ≤ 0.05, **p ≤ 0.01, ***p ≤ 0.001, and ****p ≤ 0.0001. **(C)** The expression of a few maturation markers, chemokines, cytokines in Fla^+Sia^/ds-Fla^+Sia^ matured DCs were detected by qRT-PCR. The fold change in gene expression in ds-Fla^+Sia^ vs Fla^+Sia^ matured DCs were calculated using the comparative cycle threshold (C_T_) method using the SYBR green dye. 18s rRNA was used as the housekeeping gene. For the purposes of representation, fold change of different genes in Fla^+Sia^ matured DCs is kept as 1, and the fold change in gene expression in genes of ds-Fla^+Sia^ matured DCs relative to that value is reported. The bar diagram represents data taken from at three independent experiments. Significance is represented by *p ≤ 0.05, **p ≤ 0.01, ***p ≤ 0.001, and ****p ≤ 0.0001. **(D)** The responder cells were stained with CFSE (5 µM), then incubated with Fla^+Sia^, ds-Fla^+Sia^ matured DCs (stimulators) for 7 days. These cells were further stained with anti-CD3-PE antibodies and analyzed through flow cytometry. The T cell populations were selected through PE positivity and CFSE signal in this population was assessed. T lymphocytes were positive for both CFSE and CD3-PE, and the dilution of CFSE dye indicated a higher proliferation of T lymphocytes. Representative scatter histogram is shown from one of the three indepependent experiments performed. **(E)** In a similar set up of responding, proliferating T cells, the populations of T helper (Th) and cytotoxic T cells (CTL) were identified based on CD3-PerCP, CD4-FITC and CD8-PE staining. All T lymphocytes were selected based on CD3 positivity out of which the CD4 (CD3+CD4+) and CD8 (CD3+CD8+) populations were selected. Isotype controls, single-stained controls were included for detecting populations with appropriate compensation settings. Scatter plots showing the distribution of Th and CTL populations are shown. In another setup, the responder cells were permeabilized and stained with anti FoxP3-PE antibodies for 30 min at 4°C. Cells were washed and stained with CD3-PerCP and CD4-FITC. In another setup, cells were directly stained with CD3-PerCP, CD4-FITC and CD25-APC antibodies. The Treg population present among the responder cells were detected on the basis of CD3+CD4+CD25+ as well as CD3+CD4+FoxP3+ positivity. As usual, isotype controls, single-stained controls were included for performing appropriate compensation controls. Scatter plots showing the Treg populations are shown.

The phagocytic ability of Fla^+Sia^/ds-Fla^+Sia^ matured DCs were also confirmed through confocal microscopy. The Gram negative bacteria - *Escherichia coli* ATCC 25922 were first labeled with FITC dissolved in bicarbonate buffer. The labeled bacteria were extensively washed to remove all traces of free FITC. At 37°C, the DCs were incubated for 30 min with FITC-labeled *E. coli* ATCC 25922 and then processed for confocal microscopy. The Fla^+Sia^ matured DCs internalized more bacteria (green dots in **Figure 5B**) compared to the ds-Fla^+Sia^ matured DCs. The phagocytic index, which is the average number of bacteria associated with each cell, was also calculated from the images. These observations together confirm that an increased phagocytic activity was retained by the Fla^+Sia^ matured DCs (**Figure 5B**).

In order to get a broad view of the DC maturation status, the expression of a few markers at the genetic level (mRNA) in ds-Fla^+Sia^/Fla^+Sia^ matured DCs was checked by qRT-PCR. Some of these factors were maturation markers, chemokine receptors or even cytokines. The average C_T_ values of a few genes in ds-Fla^+Sia^ and Fla^+Sia^ matured DCs were determined and normalized with respect to the housekeeping control gene (18s rRNA). The fold change in expression of each gene was calculated in respect to ds-Fla^+Sia^ vs Fla^+Sia^. For the purpose of representation, the C_T_ values of Fla^+Sia^ matured DCs have been set as 1 and the C_T_ values of ds-Fla^+Sia^ with respect of Fla^+Sia^ has been shown in **Figure 5C**. The expression of the pro-inflammatory cytokines (IFN-γ, IL-12, TNF-α) were found to be strongly upregulated in ds-Fla^+Sia^ matured DCs. Other molecules like maturation marker DC-SIGN, and chemokines (MCP-1, CCL20, CCL22) were also slightly upregulated. However, the anti-inflammatory cytokines like IL-10 and the regulatory factor - SOCS2 (suppressor of cytokine signaling 2) were slightly downregulated in ds-Fla^+Sia^ matured DCs (**Figure 5C**). Compared to Fla^+Sia^ matred DCs, the nature of ds-Fla^+Sia^ matured DCs are clearly skewed towards a more pro-inflammatory response.

The functional ability of matured DCs to stimulate T lymphocytes (CD3+) leading to their activation and clonal expansion, was assessed by T cell costimulation assay. A mixed lymphocyte reaction was set up, where responder lymphocytes (stained with CFSE dye) were incubated with flagellin matured DCs, which should result in T cell proliferation. The proliferation of responder T cells following DC stimulation could be detected by the dilution of the CFSE dye. The T cell population among the responders was also detected through CD3-PE staining **(Supplementary Figure S6)**. Responder cells incubated with unstimulated, immature DCs did not proliferate, resulting in T lymphocytes with high MFI values of CFSE dye. However, incubation with either ds-Fla^+Sia^ or Fla^+Sia^-matured DCs resulted in T cell proliferation and the dilution of CFSE dye (**Figure 5D**). Among the proliferating populations, T cells incubated with ds-Fla^+Sia^ matured DCs showed greater proliferation as reflected by the greater CFSE dye dilution leading to lower MFI values.

In a similar setup, the proliferating responder cells were stained with anti-CD3-PerCP, CD4-FITC, CD8-PE antibodies **(Supplementary Figure S6)**. The relative abundance of T helper (Th) cells (CD3+CD4+) vs cytotoxic T lymphocytes (CTLs) (CD3+CD8+) was assessed by flow cytometry. This would provide important information about the DC ability to promote T cell polarization. Stimulation with ds-Fla^+Sia^ and Fla^+Sia^ matured DCs resulted in the generation of both CTLs and Th lymphocytes. Interestingly, the ratio of cytotoxic T cells to helper T cells was higher in the case of ds-Fla^+Sia^ matured DCs than the Fla^+Sia^ matured DCs (**Figure 5E**).

DCs matured using sialylated ligands are known to promote tolerogenic immune responses which include the stimulation of T cells to differentiate into Treg cells (32, 33, 46, 47). Therefore, we have checked if there are any Treg populations within the T lymphocytes through staining with FoxP3-PE/CD4-FITC as well as CD25-APC/CD4-FITC. The responder cells (non-permeabilized) were stained with anti-CD3-PerCP, CD4-FITC, CD8-PE and CD25-APC and the population of Treg (CD3+CD4+CD25+) cells were detected (**Figure 5E**). In another setup, the responder cells were first permeabilized as per the manufacturer’s instructions and then stained with respective antibodies (CD3-V500, CD4-FITC, FoxP3-PE) (see **Supplementary Figure S6**). In both cases, the population of Treg cells as detected through CD3+CD4+CD25+ or CD3+CD4+FoxP3+ positivity, was found to be higher among the responders incubated with the Fla^+Sia^ matured DCs (**Figure 5E**).

Taken together, these observations regarding cytokine secretion, the expression of costimulatory markers, T cell polarization indicate that the application of Fla^+Sia^ resulted in the generation of matured DCs which secrete more anti-inflammatory cytokines, produces more T helper cells, and is skewed towards generating an anti-inflammatory (Th2 type) and tolerogenic response compared to the responses shown by ds-Fla^+Sia^ matured DCs. These observations are consistent with previous reports indicating that Siglecs can modulate the DC maturation process (32, 33, 46, 47).

### The status of key signaling molecules in DCs matured using flagellin

Previous results indicate that there are distinct differences in the maturation profiles of ds-Fla^+Sia^ vs Fla^+Sia^ matured DCs. We have already observed that Fla^+Sia^ bind to TLR5 and Siglec-9 on the moDC surface (**Figure 3**). Now, flagellin mediated TLR5 activation leads to a pro-inflammatory signaling cascade involving MAP kinase signaling and NF-κB activation (25, 40–43). As an inhibitory receptor, Siglec-9 activation leads to the recruitment of phosphatases-SHP1 and SHP-2 which suppress immune cell activation (17–19, 31). We suspect that the interplay between these antagonistic signaling cascades contribute to the altered DC phenotype. Accordingly, the status of a few signaling molecules which are activated downstream upon TLR5 activation, was checked in flagellin-matured human moDCs. Interestingly, the phosphorylated forms of p38 MAPK, JNK/SAPK, ERK1/2 were clearly elevated in ds-Fla^+Sia^ treated moDCs while p-Akt Ser473 and p-IκB-α showed a slight upregulation (**Figure 6A**). However, in Fla^+Sia^ matured DCs, the expression level of these phosphoproteins remained comparatively lower. This suggests that the ds-Fla^+Sia^ matured DCs show enhanced activation of TLR5 pathway compared to Fla^+Sia^ matured DCs (**Figure 6A**).

**FIGURE 6.**
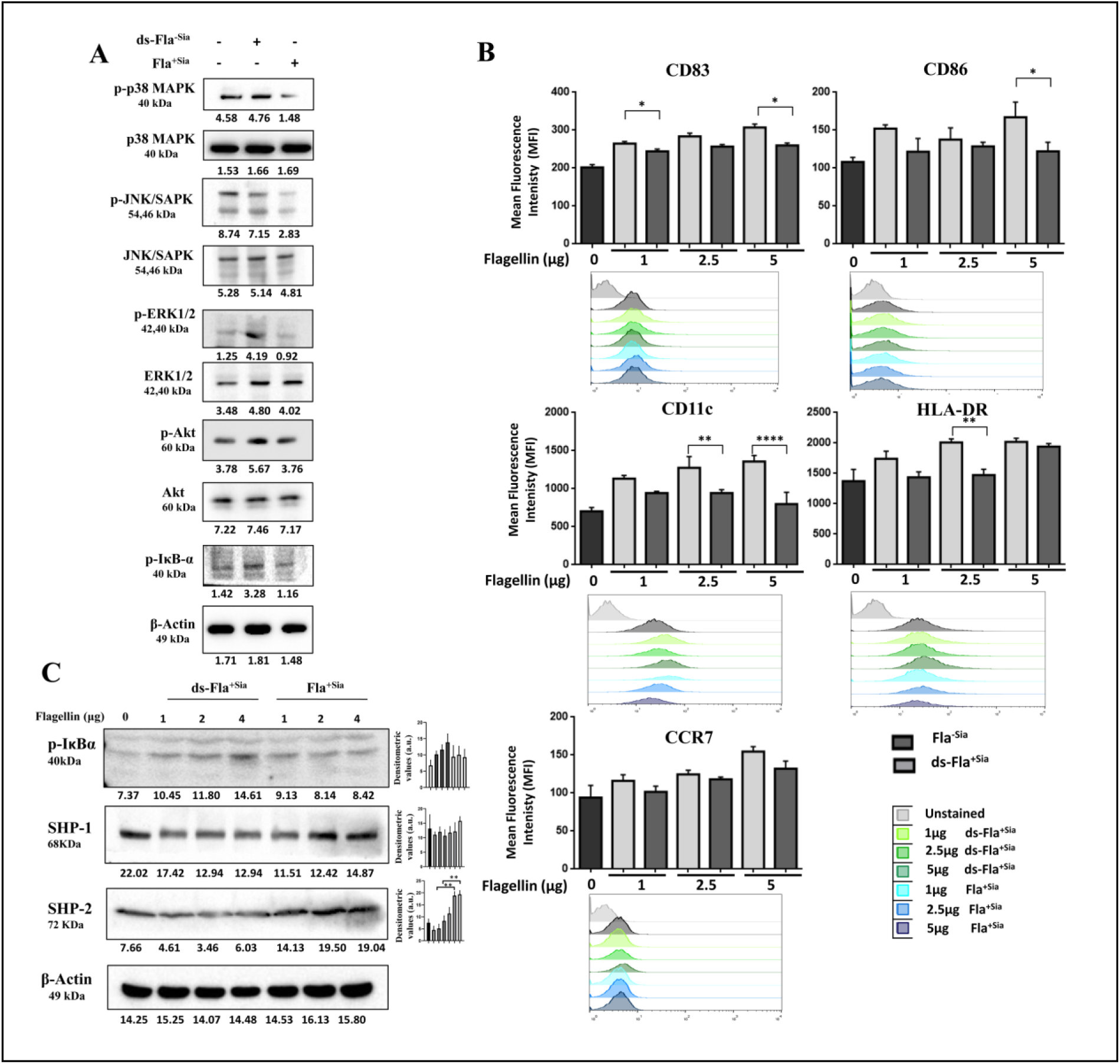
The status of TLR5 and Siglec-9 signaling in flagellin stimulated DCs. **(A)** The expression of a few important signaling molecules in moDCs was analyzed by western blotting. The cell lysates of untreated, Fla^+Sia^ and ds-Fla^+Sia^ matured DCs were resolved via SDS-PAGE, transferred to PVDF membranes and probed using primary antibodies targeting these few molecules. Blots were later incubated with appropriate secondary antibodies. Expression level of these molecules was estimated from antibody binding, which was detected using chemiluminescent substrate. The expression of β-actin was used for loading control. Experiments were independently repeated two times. **(B)** THP-1 derived DCs were stimulated with Fla^+Sia^/ds-Fla^+Sia^ (1 μg, 2.5 μg, 5 μg) as before. The status of a few costimulatory markers and maturation markers were assessed through flow cytometry. As before, untreated and treated cells were stained with cocktails of fluorophore conjugated antibodies. Unstained, single fluorophore stained controls and antibody cocktail stained samples were analyzed through flow cytometry. Appropriate compensation settings were used to visualize the populations. The data from atleast three independent experiments are depicted in the bar diagrams. Representative histograms showing shifts in stained populations are shown. Significance is represented by *p ≤ 0.05, **p ≤ 0.01, ***p ≤ 0.001, and ****p ≤ 0.0001. **(C)** The expression status of SHP-1 (Siglec-9 mediated signaling) and p-IκB-α (TLR5 mediated NF-κB activation pathway) were checked in THP-1 derived DCs following stimulation with different doses of flagellin (0, 1, 2, 4 μg). The THP-1 derived DC lysates were prepared and the expression of these proteins was probed by western blotting. Representative blots are shown while the densitometric data from three independent experiments are represented in the bar diagrams. Significance is represented by *p ≤ 0.05, **p ≤ 0.01, ***p ≤ 0.001, and ****p ≤ 0.0001.

Cell signaling studies require a large yield of matured DCs which was difficult to obtain from human moDCs. So, the status of several cell signaling molecules were explored using the human monocytic cell line - THP-1 derived DC model system (36). The DC-like cells were prepared from THP-1 cells through rhGM-CSF and rhIL-4 stimulation (see **Materials and Methods**). The THP-1 derived DCs were stimulated with increasing concentrations of ds-Fla^+Sia^/Fla^+Sia^ (1, 2.5, 5 μg) and the expression of some cell surface markers were assessed as before (**Figure 4**).

It was observed that these cells responded similarly as the moDCs to the increasing doses of Fla^+Sia^ (**Figure 6B**). The MFI values of markers – CD86, HLA-DR, CD11c – markedly remained lower in the Fla^+Sia^ treated DCs compared to the ds-Fla^+Sia^ treated DCs (**Figure 6B**). The HLA-DR expression in the 2.5 μg Fla^+Sia^ stimulated THP1-derived DCs (MFI = 1449 ± 97.3) was significantly lower compared to that observed in the 2.5 μg ds-Fla^+Sia^ stimulated THP1 cells (MFI = 1974 ± 601.7). A similar pattern was observed with CD11c (MFI = 784.7 ±79.8 vs 1353 ± 310.23), CD86 (MFI = 124 ± 49.4 vs 170 ± 99.56) in the 5μg flagellin stimulated cells. These results indicate that the THP-1 derived DCs respond in a similar fashion to sialylated flagellin as the hmoDCs. Thus, THP-1 derived DCs seem to reflect similar changes as the hmoDCs in response to flagellin, making them excellent models to explore the status of cell signaling events.

The expression of a few molecules related to the TLR5 and Siglec-9 related signaling pathways were initially assessed in THP-1 derived DCs following stimulation with 1, 2 and 4 µg of Fla^+Sia^ or ds-Fla^+Sia^. The phosphatases SHP1 and SHP2 levels were found to be upregulated upon treatment with the increasing doses of Fla^+Sia^ in the THP1-derived DCs (**Figure 6C**). This confirmed that the Siglec-9 based inhibitory signaling is predominantly active in the Fla^+Sia^ stimulated DCs. On the other hand, the levels of p-IκB-α were upregulated with the increasing doses of ds-Fla^+Sia^ treatment (**Figure 6C**). The p-IκB-α levels indicate the activation of the NF-κB pathway which is downstream of the TLR5 signaling cascade. Taken together, an enhanced activation of the TLR5 pathway and the NF-κB signaling was observed in the ds-Fla^+Sia^ matured DCs. Conversely, the inhibitory Siglec-based signaling was more active upon Fla^+Sia^ stimulation. Furthermore, the activation of Siglec-9 based inhibitory signaling in Fla^+Sia^ stimulated DCs seems to dampen TLR5 based signaling events like IκB-α phosphorylation.

### The status of TLR5, Siglec-9 signaling pathways in flagellin stimulated THP-1 derived DC following inhibitor treatment

The contribution of the TLR5 or Siglec-9 signaling pathways to DC maturation was estimated by incubating the THP-1 derived DCs with inhibitors targeting respective receptors before the cells were stimulated with flagellins. The Siglec-based inhibitory signaling requires the recruitment of phosphatases SHP-1/2 and their activity. Hence, a SHP-1/2 inhibitor - NSC-87877 was used to completely disrupt the Siglec-based immune-inhibition (48, 49). Another inhibitor-TH1020, which specifically blocks the TLR5-flagellin complex formation, was used to prevent the flagellin-based TLR5 activation (50).

The activation of TLR5 or Siglec signaling leads to the recruitment of their downstream signaling partners to respective receptors. This association was evaluated in flagellin stimulated THP-1 derived DCs. TLR5 associates with a few factors like MyD88, TRAF6, IRAK-4 for signal transduction while Siglec-9 recruits the phosphatases SHP1 and SHP-2. To measure these associations, antibodies targeting TLR5 and SHP-1were used to set up immunoprepicitations. THP-1 derived DCs were singly treated with (2 µg) Fla^+Sia^ or ds-Fla^+Sia^. Some DCs were pretreated with either NSC-87877 (to block Siglec signaling) or with TH1020 (to block TLR5 signaling) followed by the stimulation with Fla^+Sia^. The cell lysates prepared from these treated cells were incubated with TLR5 or SHP-1 targeting antibodies and then later incubated with Protein A-agarose beads. The proteins which remained bound to the beads were precipitated, then resolved via SDS-PAGE and analyzed by western blotting. An increased association between TLR5 with MyD88, TRAF6 and IRAK-4 was observed in ds-Fla^+Sia^ treated DCs (**Figure 7A**), while this association remained low in Fla^+Sia^ treated DCs. Interestingly, inhibiting the SHP-1/2 activity in Fla^+Sia^ treated cells with NSC-87877 resulted in an increase in TLR5-MyD88/TRAF6/IRAK-4 association. However, the blocking of flagellin binding with TLR5 by TH1020 nearly completely abolished the association between TLR5 with TRAF6, and IRAK4 (**Figure 7A**) in Fla^+Sia^ treated DCs.

**FIGURE 7.**
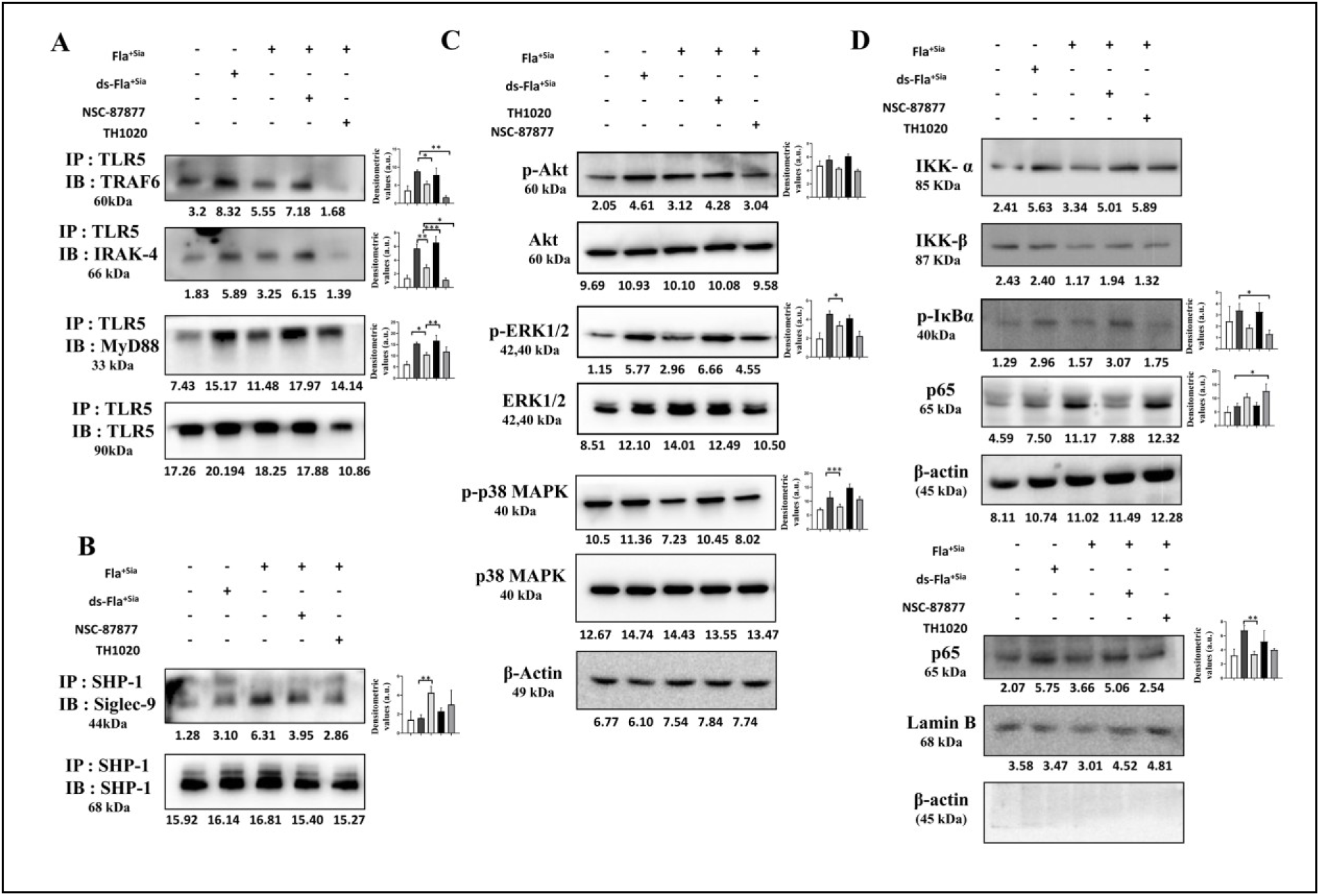
Inhibitor based modulation of TLR5 and Siglec-9 signaling pathways in flagellin stimulated THP-1 derived DCs. **(A)** Flagellin treated THP-1 derived DCs were treated with a SHP-1/2 inhibitor (NSC-87877) or flagellin-TLR5 complex inhibitor (TH1020) in order to disrupt Siglec-9 and TLR5 signaling respectively. Cells were then treated with Fla^+Sia^ or ds-Fla^+Sia^ (2 µg). After 24 hr treatment, cell lysates were prepared and the association of TLR5 with other downstream interacting molecules was examined by co-immunoprecipitation. Briefly, lysates were overnight incubated with anti TLR5 antibodies at 4°C. Next day, the mix was incubated with protein A-agarose beads for 3 hours at 4°C. The beads were washed and all remaining bead-bound proteins were extracted into sample buffer. The proteins were resolved by SDS-PAGE, transferred to PVDF blots and porbed by western blotting. **(B)** Following similar treatment, the association between Siglec-9 and SHP-1 was probed by co-immunoprecipitation. Lysates were incubated with anti SHP-1 antibodies for overnight at 4°C. The antibody bound proteins were separated through incubation with protein A-agarose. The association between SHP-1 and other proteins were later probed by western blotting. **(C)** The status of several phosphorylated downstream signaling molecules involved in the TLR5 signaling cascade was checked in cell lysates prepared from the treated, flagellin stimulated DCs by western blotting. The flagellin treated THP-1 derived DCs were separated into their cytosolic and nuclear fractions. The signaling molecules involved in the NF-κB pathway as well as the nuclear translocation of NF-κB p65 subunit was assessed by western blotting.

In all cases, representative blots showing the change in protein expression are selected from the at least three independent experiments performed. The densitometric data collected from these experiments are represented in the histograms. Significance is represented by *p ≤ 0.05, **p ≤ 0.01, ***p ≤ 0.001, and ****p ≤ 0.0001.

On the other hand, the association between Siglec-9 and SHP-1 was observed to be highest in Fla^+Sia^ treated DCs (**Figure 7B**). In case of pretreatment with NSC-87877, TH1020 followed by Fla^+Sia^ treatment, a reduction in Siglec-9-SHP-1 association was observed. As expected, the Siglec-9-SHP-1 association was nearly absent in ds-Fla^+Sia^ stimulated DCs. These observations confirm that stimulation with sialylated flagellin leads to specific activation of Siglec-9 signaling and increased the Siglec-9-SHP-1 association. Interestingly, this is accompanied by the suppression of the flagellin-based TLR5 activation and TLR5 association with MyD88, TRAF6 and IRAK-4 is reduced. These results suggest that Siglec-9 activation directly interferes with the TLR5-flagellin based signaling.

The expression of a few important downstream signaling molecules was also checked. An increased expression of phosphoproteins like p-ERK1/2 and p-p38-MAPK was observed in ds-Fla^+Sia^ treated DCs possibly due to enhanced TLR5 signaling. Similar enhanced expression was also seen in Fla^+Sia^ treated DCs when the Siglec-based inhibitory signaling was suppressed by NSC-87877 **(Figure 7C)**. However, the levels of these phosphoproteins remained low in cells where Siglec signaling was dominant. This was observed in cells treated with only Fla^+Sia^ or with Fla^+Sia^ along with the TLR5-flagellin complex blocker (TH1020) (**Figure 7C**).

TLR5 activation leads to a signaling cascade that culminates in the nuclear translocation of the NF-κB p65 subunit. The cytosolic and nuclear fractions of flagellin-treated cells were separated and the status of a few proteins related to the NF-κB signaling was probed. The ds-Fla^+Sia^ treated DCs showed an increase in IKK-α, IKK-β, p-IκB-α levels in the cytosol while the NF-κB p65 subunit level in the cytosol was reduced (**Figure 7D**). An opposite scenario was observed in the DCs treated with Fla^+Sia^. Lowered levels of IKK-α/β and p-IκB-α were observed in the cytosol while the level of p65 subunit was greater than that seen in ds-Fla^+Sia^ stimulated DCs. In the nuclear fraction of ds-Fla^+Sia^ stimulated DCs, the signal from the p65 subunit was much greater than that seen in Fla^+Sia^ stimulated DCs (**Figure 7D**). This indicates that the NF-κB pathway is activated more upon ds-Fla^+Sia^ stimulation, which enhances the nuclear translocation of the p65 subunit.

In the case of NSC-87877 treatment, the amount of IKK-α and p-IκB-α was slightly upregulated while there was a reduction in the level of cytosolic p65 (**Figure 7D**). Again, TH1020 treatment resulted in a similar scenario as DCs treated with only Fla^+Sia^. In the nuclear fractions, NSC-87877 treated lysates showed more signal of the p65 subunit while this signal was nearly abolished upon blocking flagellin-TLR5 interaction through TH1020 treatment. The absence of any β-actin bands in the nuclear fraction indicates that the nuclear fractions did not contain too much of cytosolic contaminants (**Figure 7D**).

### The status of TLR5, Siglec-9 signaling pathways in THP-1 derived DCs after siRNA treatment

In order to gain a better understanding of the interplay between Siglec-9 and TLR5 signaling pathways in case of Fla^+Sia^ stimulation, the expression of TLR5 and Siglec-9 in THP-1 derived DCs were suppressed through RNA interference using siRNAs. Initially, the DCs were transiently transfected with commercially purchased siRNA and cell lysates were prepared. As a control, no siRNA were added in the case of the mock transfectants. Western blotting confirmed a reduction in the TLR5, Siglec-9 protein levels in these siRNA transfected cells (**Figure 8A**). Additionally, the mock transfectant and siRNA transfectant cells were stained with anti-TLR5 and anti-Siglec-9 antibodies and PE-tagged secondary antibodies followed by flow cytometry. At 24 hr post transfection, flow cytometry revealed a reduction in the expression of TLR5 and Siglec-9 on the siRNA transfected cell surfaces (**Figure 8B**).

**FIGURE 8.**
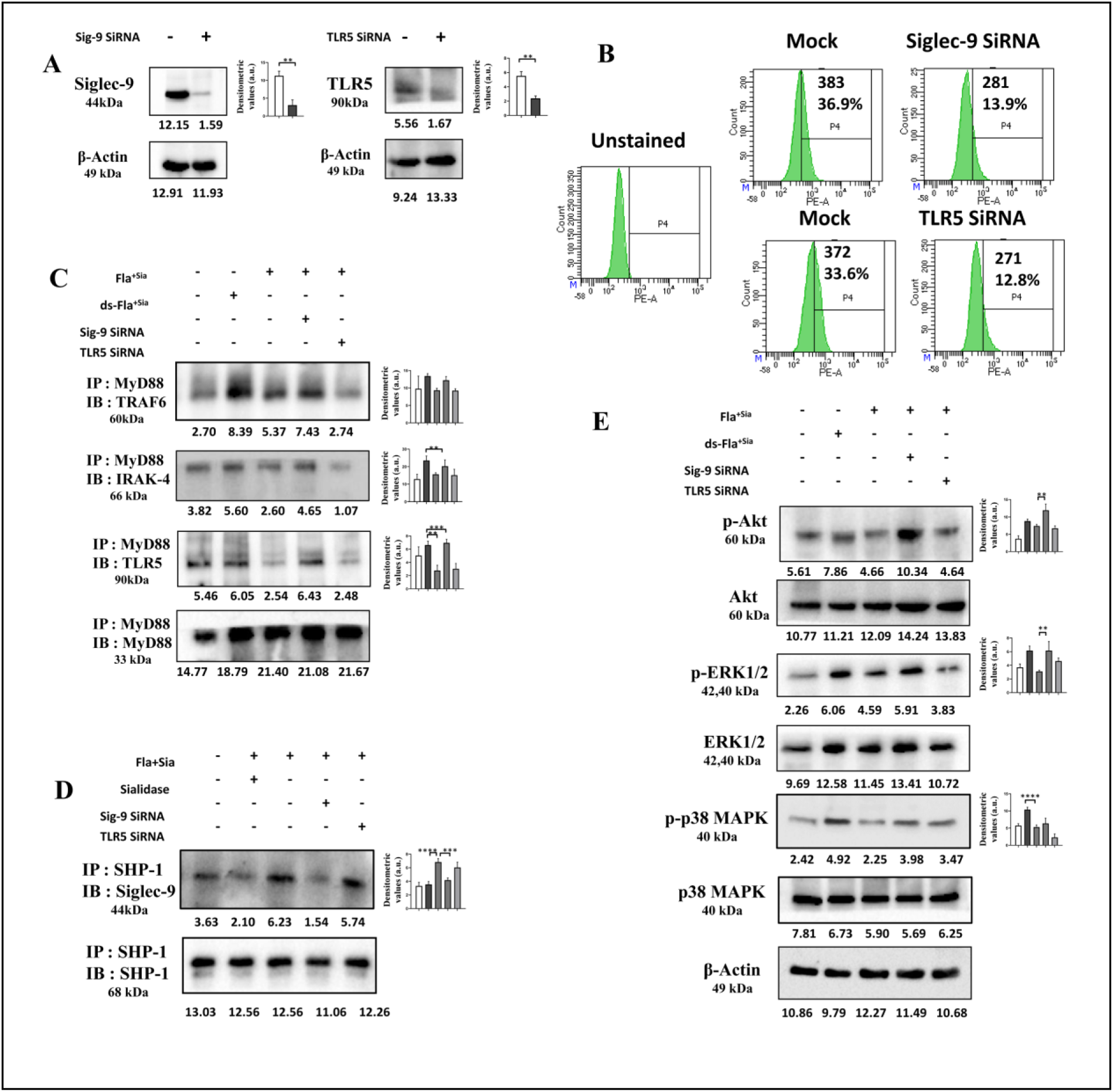
The status of cellular signaling in flagellin stimulated THP-1 derived DCs following siRNA transfection. **(A)** A siRNA based transient knockdown of Siglec-9 and TLR5 expression was performed. Following 24 hr of transfection, cells were lysed and then analyzed by western blotting. The expression levels of Siglec-9 and TLR5 were compared between transfected and non-transfected cells. A representative blot out of three independent experiments is shown here. **(B)** As before, siRNA transfected cells were stained with anti Siglec-9 and anti TLR5 antibodies followed by counterstaining with PE tagged secondary antibodies. The expression of Siglec-9, TLR5 expression on cell surface was then detected by flow cytometry. **(C)** Non-transfected and siRNA transfected DCs were treated with Fla^+Sia^/ds-Fla^+Sia^ as before. Cell lysates were prepared and lysates were set up for co-immunoprecipitation using anti MyD88 antibodies. Antibody bound proteins were precipitated using protein A-agarose beads. The association between MyD88 with TLR5, TRAF6 and IRAK-4 were examined by western blotting. **(D)** In a similar set up, DC lysates were prepared and set up for co-immunoprecipitation using the anti SHP-1 antibodies. The association between Siglec-9 and SHP-1 was assessed in transfected vs non-transfected cells through western bloltting. **(E)** The status of several phosphorylated pathway molecules involved in TLR5 signaling was also compared in the cell lysates of transfected vs non-transfected cells by western blotting. In all the cases of western blotting, representative blots showing the change in protein expression are selected from one of the at least three independent experiments performed. The densitometric data collected from these experiments are represented in the histograms. Significance is represented by *p ≤ 0.05, **p ≤ 0.01, ***p ≤ 0.001, and ****p ≤ 0.0001.

The status of the TLR5 and Siglec-9 pathways was again evaluated in THP-1 derived DCs. Cells were transfected with siRNA for 24 hr to allow a transient knockdown of TLR5 or Siglec-9 expression followed by a further 24 hr stimulation with Fla^+Sia^. Control cells were stimulated with Fla^+Sia^ or ds-Fla^+Sia^. As before, the association between TLR5 and its adapter proteins and that between Siglec-9 and its downstream partners – SHP1/2 were probed through co-immunoprecipitation. Since, the TLR5 and Siglec-9 expression is being modulated here; immunoprecipitations were performed using anti-MyD88 and anti-SHP-1 antibodies.

The association between MyD88 and TRAF6, IRAK-4 was markedly higher in ds-Fla^+Sia^ treated DCs as compared to Fla^+Sia^ treated cells (**Figure 8C**). This corroborates with all previous observations that TLR5 signaling is more active in ds-Fla^+Sia^ stimulated cells. In case of Siglec-9 silencing, the association between MyD88 and TRAF6, IRAK-4 and TLR5 remained higher than that seen with only Fla^+Sia^ stimulated DCs. However, the silencing of TLR5 resulted in a reduction of all MyD88-TRAF6-IRAK-4-TLR5 association. In summary, when Siglec-9 signaling was active, as seen in only Fla^+Sia^ stimulated DCs, the TLR5-MyD88 association remained low. Interestingly, the knockdown of Siglec-9 enhanced the association of TLR5 with MyD88. Comparing the binding results from DCs stimulated with only Fla^+Sia^ and Siglec-9 silenced DCs stimulated with Fla^+Sia^; we can infer that the interactions between flagellin sialic acids and Siglec-9 inhibits the association of TLR5 with its downstream adapter molecules (**Figure 8C**). A reversed scenario was observed in the case of SHP-1-Siglec-9 association. The Fla^+Sia^ treated DCs showed a greater association between SHP-1 with Siglec-9 unlike ds-Fla^+Sia^ treated cells. In the Fla^+Sia^ treated cells, this SHP-1-Siglec-9 association remained high when TLR5 was silenced but was abolished in the case of Siglec-9 knockdown (**Figure 8D**).

In summary, TLR5 signaling - represented by the association between TLR5-MyD88, TRAF6, IRAK4; was enhanced in ds-Fla^+Sia^ treated cells and in Fla^+Sia^ treated cells when Siglec-9 was silenced. However, only Fla^+Sia^ treated cells exhibit an increased activation of the Siglec-9 inhibitory signaling pathway, shown by an increased association between Siglec-9-SHP-1. This Siglec-9-SHP1 interaction may have possibly suppressed the activation of the TLR5 pathway, leading to the lower association between TLR5-MyD88, IRAK-4, TRAF6.

As before, the expression patterns of some of the secondary phosphorylated signaling molecules were also checked. As observed before, the levels of p-AKT, p-ERK1/2, p-P38 MAPK remained enhanced in ds-Fla^+Sia^ treated DCs compared to Fla^+Sia^ treated DCs. However, the silencing of TLR5 is accompanied by a reduction in the expression levels of the same signaling molecules. This again confirmed that TLR5 based detection of PA flagellin is critical for the generation of a strong pro-inflammatory signaling cascade (**Figure 8E**).

Interestingly, a transient knockdown of Siglec-9 expression followed by Fla^+Sia^ treatment resulted in an elevation in the levels of these phosphoproteins (**Figure 8E)**. Taken together, these observations confirm that Siglec-9 exerts a strong immunosuppressive action on the TLR5 based cell activatory signaling.

### The impact of modulating TLR5, Siglec-9 signaling pathways on DC maturation process

Our observations till now indicate that the Siglec signaling suppresses TLR5 based immune-activatory signaling and reduces the levels of several downstream signaling molecules. We propose that these cellular signaling events caused by the antagonistic pathways lead to the different phenotypes and functional abilities exhibited by ds-Fla^+Sia^/Fla^+Sia^ matured DCs (**Figure 4, 5**). We have now explored if the effect of modulating Siglec-9 and TLR5 expression has any effect on DC functions like cytokine responses and the NF-κB pathway activation.

The status of the NF-κB pathway following flagellin stimulation was directly measured using a NF-κB p65 promoter-driven luciferase probe. Activation of the NF-κB pathway at the downstream of the TLR5-driven signaling cascade results in the release of the p65 subunit, which is translocated into the cell nucleus. Here, free p65 subunit can bind to the promoters of this NF-κB p65 probe, which leads to the synthesis of the luciferase enzyme. Hence, the quantitation of luciferase production will give an estimate of the NF-κB pathway activation. The THP-1 derived DCs were transfected with this probe and then were treated with inhibitors – NSC-87877 or TH1020. In other cases, the DCs were co-transfected with TLR5, Siglec-9 targeting siRNA and the NF-κB probe. In all cases, these DCs were stimulated with Fla^+Sia^/ds-Fla^+Sia^ for 24 hours. Luciferase production, indicating NF-κB activity was measured as per the manufacturer’s instructions **(Figure 9A)**. As observed before, NF-κB activation was observed to be higher in ds-Fla^+Sia^ treated DCs compared to Fla^+Sia^ treated DCs, indicating greater activation of the TLR5 pathway. Targeting the Siglec-9 receptor through siRNA or SHP1/2 using inhibitors respectively resulted in a similar increase in NF-κB activation even under Fla^+Sia^ stimulation. When the TLR5-flagellin association was blocked using TH1020 or through siRNA based TLR5 silencing, the NF-κB activation was markedly reduced **(Figure 9A)**.

**Figure 9.**
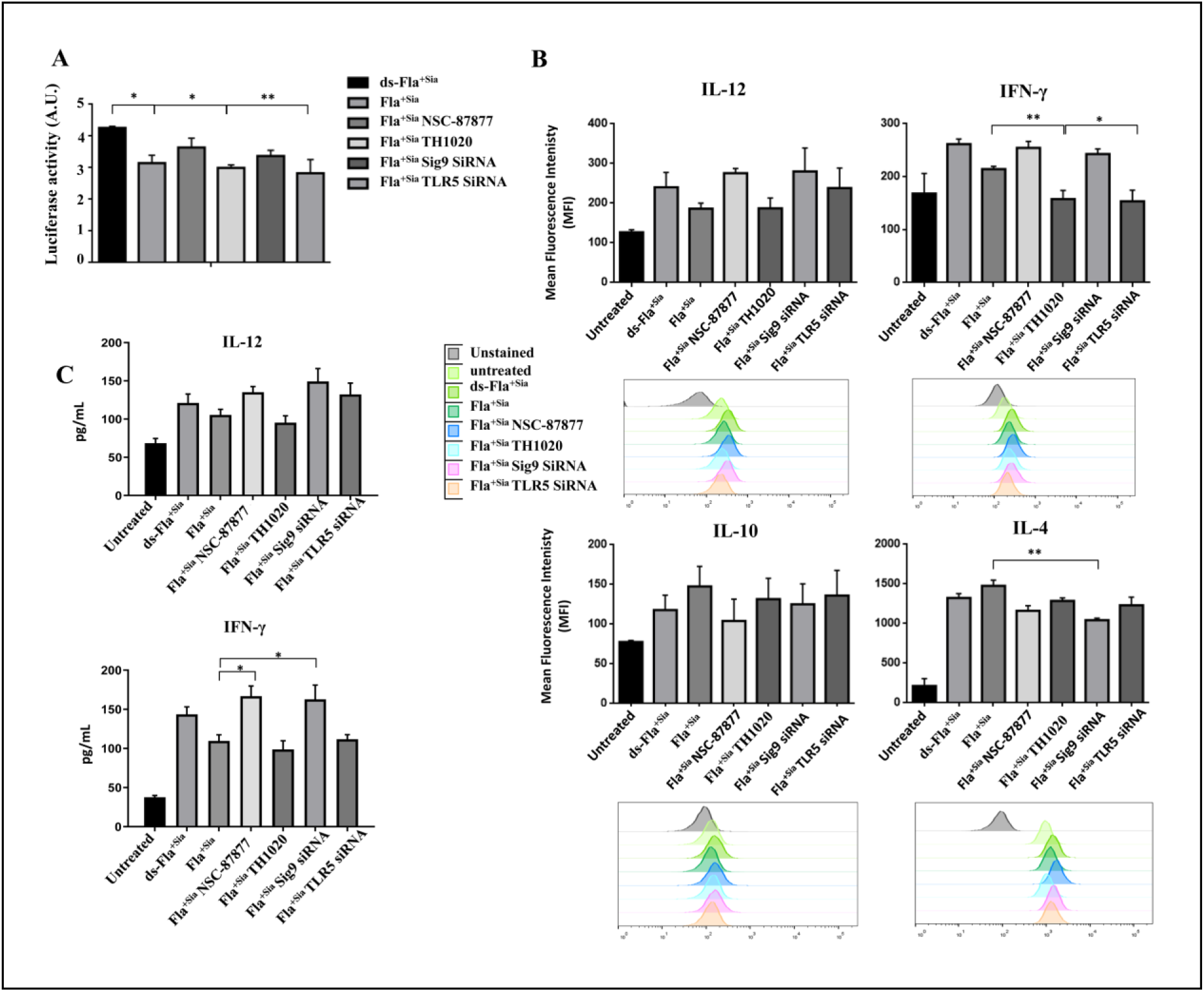
The effect of Siglec-9 mediated suppression of TLR5 signaling on cellular response by flagellin stimulated DCs. **(A)** The activation of NF-κB pathway was evaluated using a NF-κB reporter plasmid which was transfected into the THP1-derived DCs. In order to disrupt TLR5/Siglec-9 signaling, cells were also contransfected with TLR5/Siglec-9 targeting siRNA or were treated with NSC-87877 or TH1020. Cells were then stimulated with Fla^+Sia^ or ds-Fla^+Sia^ (2 µg) overnight and then lysates were prepared. The production of the luciferase enzyme in the lysates was detected following the manufacturer’s instructions. Readings were taken in a plate multireader capable to detecting luminescence. The bar diagram is constructed from the data of three independent experiments. Significance is represented by *p ≤ 0.05, **p ≤ 0.01, ***p ≤ 0.001, and ****p ≤ 0.0001. **(B)** As before, Flagellin treated DCs with siRNA or inhibitor based suppression of TLR5 and Siglec-9 signalling was analyzed by flow cytometry to detect the intracellular levels of cytokines produced. As before, untreated and treated cells were stained with cocktails of fluorophore conjugated antibodies. Unstained, single fluorophore stained controls and stained samples were analyzed through flow cytometry. Appropriate compensation settings were used to visualize the populations. The data from atleast three independent experiments are depicted in the bar diagrams. Representative histograms showing shifts in stained populations are shown. Significance is represented by *p ≤ 0.05, **p ≤ 0.01, ***p ≤ 0.001, and ****p ≤ 0.0001. **(C)** DCs transfected with siRNAs targeting TLR5/Siglec-9 or treated with TH1020/NSC-87877 were overnight stimulated with Fla^+Sia^. The culture supernatnats were collected and stored at-80°C. Cytokine levels in this supernatant were detected by using IL-10, IL-4, IFN-γ ELISA kits following the manufacturer’s instructions.

The NF-κB pathway plays a critical role in the DC maturation process (51) and has a profound effect on the cytokine response. So, the THP-1 derived DCs treated with inhibitors/siRNA were stimulated with Fla^+Sia^/ds-Fla^+Sia^ and then cytokine response was evaluated by flow cytometry. When Fla^+Sia^ was used as a stimulus, the blocking of Siglec-based inhibitory signaling by inhibiting SHP-1/2 activity via NSC-87877 resulted in the increased production of pro-inflammatory cytokines like IFN-γ and IL-12 **(Figure 9B)**. Upon siRNA based knockdown of Siglec-9, similar results were obtained **(Figure 9B)**. Interestingly, this response was similar to the DC cytokine response against ds-Fla^+Sia^. The inhibitor (TH1020) based disruption of the TLR5-flagellin interaction resulted in a reduction of levels of these same proinflammatory cytokines **(Figure 9B)**. Again, the silencing of TLR5 also negatively affected the level of IFN-γ cytokines in Fla^+Sia^ treated DCs.

The production of anti-inflammatory cytokines (IL-4, IL-10) followed an opposite trend. Compared to only Fla^-Sia^/Fla^+Sia^ treated DCs, SHP-1/2 inhibitors lead to a reduction in cytokine production. Blocking TLR5-flagellin association with TH1020 or by silencing TLR5 by siRNA resulted in slight increases in anti-inflammatory cytokine production (IL-10, IL-4) in response to Fla^+Sia^ treatment **(Figure 9B)**.

The observed trend in cytokine production was also confirmed by ELISA. Cells where TLR5/Siglec-9 signaling (via inhibitors or siRNA transfection) was disrupted were stimulated with Fla^+Sia^ as before. As a control, untreated cells and normal cells stimulated with Fla^+Sia^ or ds-Fla^+Sia^ were also kept. Culture supernatants from all these cells were collected and the levels of IFN-γ, IL-12 were estimated. The observations indicate that DCs produce more pro-inflammatory cytokines in response to ds-Fla^+Sia^ than Fla^+Sia^, which was also previously observed (**Figure 4B**). Targeting the TLR5 signaling resulted in a lowered secretion of these cytokines **(Figure 9C)**. However, when Siglec-9 signaling is disrupted (siRNA/NSC-87877), the level of these cytokines remained elevated **(Figure 9C)**. In brief, the cytokine response by flagellin stimulated DCs clearly demonstrate that TLR5 signaling is important in maintaining the proinflammatory response by the DCs. It also shows that the Siglec-9 signaling alters such DC responses, so that the inhibtition of this pathway allows DCs to respond with higher secretion of IFN-γ, IL-12.

Taken together, the inhibition of Siglec-9 and associated SHP1/2 activity results in an increase in TLR5 activation, signaling. This in turn promotes NF-κB pathway activation such that the matured DCs produced show a strong pro-inflammatory cytokine response. Through its immunosuppressive action, Siglec-9 modulates this entire process and the resulting DC phenotype. Thus, the suppression of Siglec-9 signaling allows more NF-κB activation, such that even the Fla^+Sia^ stimulated DCs show a higher secretion of pro-inflammatory cytokines. Therefore, the interplay of the TLR5, Siglec-9 pathways in DCs matured with Fla^+Sia^/ds-Fla^+Sia^ determines the outcome of DC maturation.

## Discussion

The presence of sialic acids in PA and its role in modulating neutrophil and macrophage responses against the bacteria has been previously reported (4, 5, 34). PA sialic acids interact with the host immune cell surface Siglecs to initiate a Siglec-based inhibitory signaling which in turn suppresses anti-bacterial responses (5, 34). Flagellin was one of the several sialoglycoproteins purified and identified from PA^+Sia^ (6). Here, we have explored the sialylation status of flagellin and its role in modulating the process of DC maturation. The presence of sialic acids in flagellins isolated from PA^+Sia^ (Fla^+Sia^) has been confirmed through biochemical assays, lectins and Siglecs binding. Fla^+Sia^ were found to interact with both TLR5 and Siglec-9 receptors present on the immature monocyte-derived dendritic cells (moDCs). Interestingly, the DCs matured using either sialylated (Fla^+Sia^) or desialylated flagellins (ds-Fla^+Sia^) showed slightly different maturation profiles. These cells differed in the expression of surface markers, pro-inflammatory cytokines secretion, T cell polarization and showed altered cell signaling events. Generally, the TLR5-based detection of flagellin leads to immune cell activation represented by robust MAP Kinase signaling, NF-κB activation and pro-inflammatory cytokines secretion. Here, the additional interactions between Siglec-9 and Fla^+Sia^ resulted in the activation of SHP1 phosphatase-based inhibitory signaling. This signaling axis suppressed the pro-inflammatory TLR5 signaling pathway. The silencing of Siglec-9 or the use of SHP-1/2 phosphatase inhibitors resulted in an increased TLR5-MyD88 association which led to an enhanced MAPK signaling, NF-κB nuclear translocation and secretion of pro-inflammatory cytokines by Fla^+Sia^ matured DCs. These observations indicate that the PA sialylated flagellins can modulate the process of DC maturation through the engagement of cell surface Siglec-9, which then interferes with the TLR5 pathway and leads to an altered DC maturation profile (**Figure 7**).

Flagellin is a highly conserved protein which polymerizes to form the flagellar filament. The PA flagellins are classified into two distinct types-a or b, based on their reactions with antiserum. The PA strains like PAO1, PA14 produce the b-type flagellin while several clinical isolates and the well-known PAK strain possess the a-type flagellin (9, 10). These two serotypes share 70% identical nucleotide sequences along with some variations in their sequence and glycosylation. The b-type flagellin are more conserved across strains and encode a fixed 53 kDa flagellin protein while the a-type flagellins vary in their molecular weights (45-52 kDa) due to extensive posttranslational modifications. In the PAK strain, a large flagellar glycosylation island of 14 genes controls the glycosylation of the a-type flagellin (9). In contrast, the b-type flagellins are not extensively glycosylated, with only four genes being identified in the flagellar glycosylation locus of the PAO1 strain (10). The b type flagellin was considered to lack any glycosylation till a small glycan moiety (709 Da) comprising of a single deoxyhexose monosaccharide attached to a phosphate group was identified in the strain PAO1. This moiety is attached via O-glycosylation at Ser 191/Ser 195 of the b-type flagellin (10). Another recent study has identified that the b-type flagellin additionally contains an N, N-dimethylthreonine residue (52). A similar flagellin glycosylation pattern is expected in strain PA14, since it shares 99% identical genome sequence with PAO1. A later study identified additional three N-linked glycosylation sites at Asn26/Asn69/Asn439 of the PA14 flagellin (53). Thus, the full extent of glycosylation present in the b-type flagellin of PA14 has not been completely deciphered and it may be more complex than previously thought. PA flagellin has been described to have a potential use as vaccines (7, 37, 54–57) and flagellin glycosylation is also important for its immune-protective role (58, 59). In this respect, we have explored the sialylation status of flagellins from PA14 grown in the presence of FCS, and explored the impact of such sialylation in modulating immune cell function. Initially, we have used biochemical assays, lectin and Siglec binding to show the presence of sialic acid residues in the b-type flagellin of PA14 (**Figure 1 and 2**). More importantly, sialidase treatment of the isolated flagellin protein abrogated such lectin, Siglec binding - providing further proof of the presence of sialic acids (**Figure 2**). Although the mechanisms of sialic acid uptake by PA are still unknown, the presence of sialic acids and pseudaminic acids has been previously reported in PA (4–6, 60). Deciphering the sialoglycan attachment site and elucidating the complete structure of the flagellin sialoglycan is the next logical step in our study of the PA14 flagellin.

**Figure 7.**
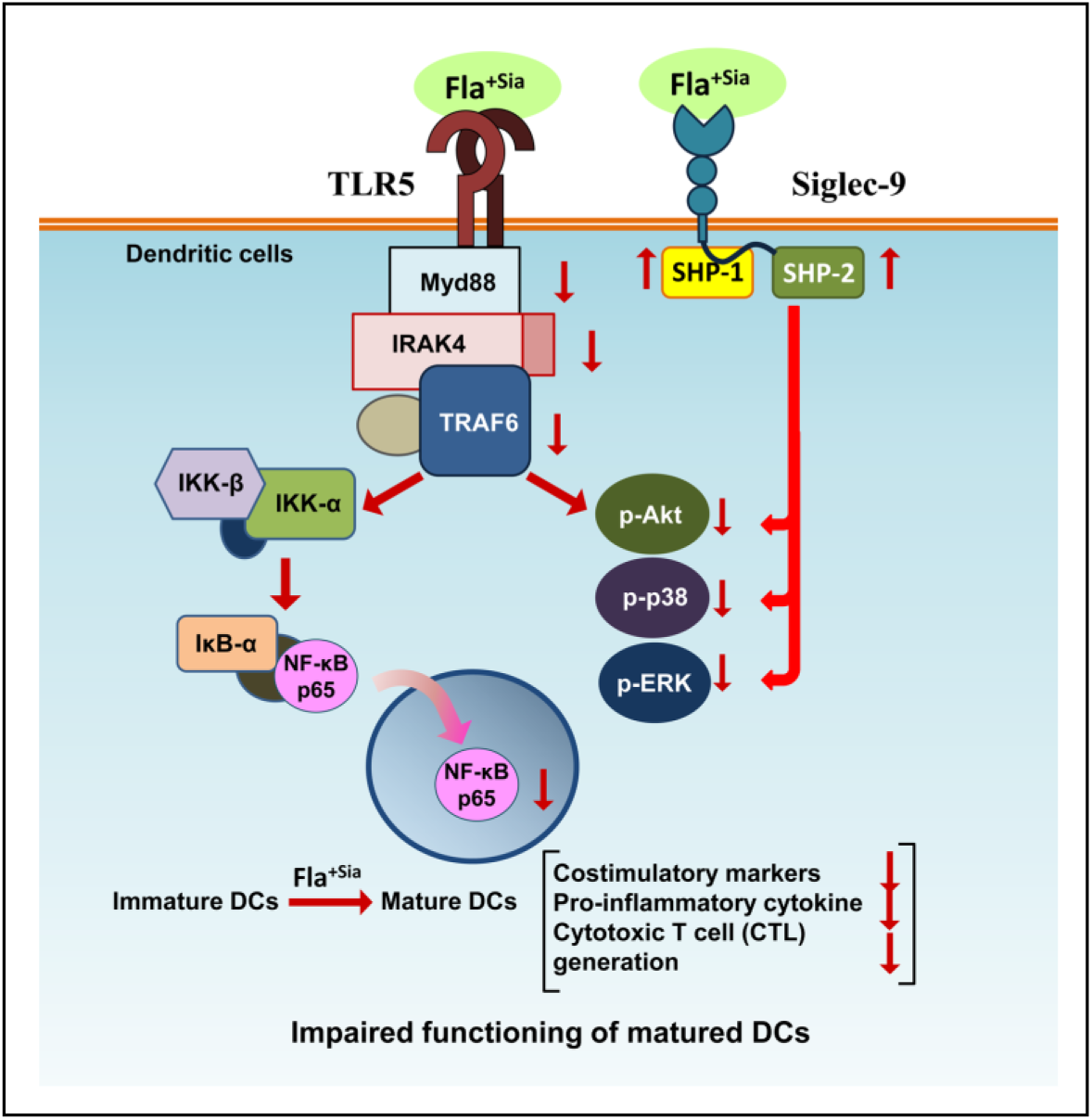
A schematic diagram representing the modulation of signaling pathways in Fla^+Sia^ stimulated DCs through multiple interactions involving the receptors-TLR5 and Siglec-9.

Dendritic cells are an important group of immune cells which are potent antigen presenting cells (APCs) responsible for the capture, processing and presentation of antigens via the Major Histocompatibility Complexes (MHCs). Additionally, DCs express several costimulatory molecules which can interact with their cognate receptors on other immune cells, thereby activating them through costimulation (19). DCs function as phagocytic APCs while they can also induce B and T cell activation and proliferation. Therefore, DCs mediate a link between the innate and adaptive arms of the immune system. The process of DC maturation and the factors involved in it determine the nature and function of the resulting matured DCs (23). In turn, through their costimulatory activity, DCs regulate the type of T cells (Th1/Th2 or Treg) that will be generated. Thus, DCs play an important role in determining the nature and vigor of immune responses. Several bacterial pathogens are known to target DCs considering their significant role in the immune system. The pathogen *Campylobacter jejuni* targets DC maturation and DC-mediated T cell polarization (15). *Brucella abortus* regulates DC maturation by modulating the TLR signaling pathways (61, 62) while *Helicobacter pylori* infection arrests DCs at a “semi-mature state” (63, 64). PA is also known to modulate DC activity through the quorum-sensing molecules, which represses IL-12 production by the LPS matured murine bone marrow-derived DCs (bmDCs) and impedes the DC based T-cell proliferation (65). Here, we provide evidence that the sialylated flagellin is another PA virulence factor which can suppress DC activity by interfering with the process of DC maturation (**Figure 7**).

Generally, flagellin monomers can be rapidly detected by the host cell surface TLR5 (66). The TLR5 is expressed by nearly all immune cells as well as epithelial cells. As a pattern recognition receptor (PRR), it is capable of detecting flagellin from nearly all types of motile bacteria. It is widely expressed and TLR5 orthologues have been identified in mammals and fishes (66, 67). Flagellin is the only identified ligand for TLR5 and its detection leads to the induction of a strong pro-inflammatory response. Due to its high immunogenicity, flagellin can be used as a maturing stimulus to generate matured DCs (25, 41–43). Here, we again confirm that PA flagellin is able to induce DC maturation (**Figure 4**). However, DCs matured using sialylated (Fla^+Sia^) or desialylated flagellin (ds-Fla^+Sia^) vary slightly in their maturation profiles (**Figure 4 and 5**). Furthermore, these cells exhibit signficant differences in cell signaling events (**Figure 6**).

Here, we have established that the Fla^+Sia^ binds to moDC surface through the receptors TLR5 as well as Siglec-9 (**Figure 3**). Interestingly, Siglec-9 is a sialic acid binding immunoglobulin-type receptor which contains an immunoreceptor tyrosine based inhibitory motifs (ITIMs) in its cytoplasmic tail (19). Siglec-9 interacts with cell surface (cis interaction) or extracellular sialylated ligands (trans interaction) and promotes immunosuppression through the recruitment of SHP-1/2 phosphatases (18). Accordingly, we have observed that ds-Fla^+Sia^ treated DCs show an increased activation of TLR5 pathway and an enhanced association between TLR5 with MyD88/TRAF6/IRAK-4 (**Figure 7 and 8**). Consequently, an enhanced activation of MAP kinase related signaling molecules (Akt, ERK1/2, p38-MAPK) and an increased nuclear translocation of NF-κB p65 subunit was observed. However, these observations were reversed in Fla^+Sia^ treated DCs, where an increased association between Siglec-9 and SHP-1 was observed. We propose that the altered phenotypes of ds-Fla^+Sia^/Fla^+Sia^ matured DCs were due to differences in cellular signaling events. We observed that disrupting Siglec-9 signaling by siRNA based knockdown or inhibitor based suppression of SHP1/2 activity, led to an enhanced TLR5 signaling in Fla^+Sia^ stimulated DCs. Consequently, the DC responses towards Fla^+Sia^ were reversed. Furthermore, DCs matured using Fla^+Sia^ secrete more pro-inflammatory cytokines when Siglec-9/SHP-1 signaling has been disrupted (**Figure 6**). Taken together, these observations indicate that the additional interaction between Siglec-9-sialic acids of Fla^+Sia^ negatively impacts the process of DC maturation. The Siglec-9-Fla^+Sia^ interactions generate immune-inhibitory signaling through SHP1/2, which suppresses the pro-inflammatory signaling generated from TLR5-flagellin interactions. The TLR5 mediated activation of kinase-based signaling and NF-κB pathways are generally inhibited by the dephosphorylating activities of SHP1/2 phosphatases. Interestingly, we have observed here that the TLR5 association with MyD88/TRAF6/IRAK-4 was also reduced in Fla^+Sia^ stimulated DCs, which could be reversed upon suppressing Siglec-9 signaling. Among the protein factors involved in these interactions, only SHP1 is known to directly bind and interact with MyD88 in IL-1 signaling (68, 69). Similarly, Siglec-E (murine ortholog of Siglec-9) based direct modulation of the sialylated receptor TLR4 is also known (70–72). However, TLR5 has not yet been identified to be sialylated and no direct interactions between Siglecs and TLR5 have been yet reported. Thus, our observation that Siglec-9 signaling lowered TLR5 association with MyD88/TRAF6/IRAK-4 might not be due to direct interactions between any of these factors. Instead, this observation may be possibly explained by the fact that both TLR5 and Siglec-9 compete to bind with the same ligand (Fla^+Sia^) on DC surfaces (**Figure 7**). This binding competition may result in lower TLR5 binding and activation when Siglec-9 is also active in interacting, binding with Fla^+Sia^. Interestingly, the addition of sialic acids to flagellin leading to a lowered TLR5 activation may be an effective strategy to suppress the activation of cells which encounter PA flagellins.

The involvement of inhibitory receptors during DC maturation is known to directly influence the DC maturation process, which affects DC activities like T cell polarization. Several studies have investigated the inhibitory and modulatory role of sialoglycans and Siglecs on DC activity. Hypersialylated ligands from oncoproteins found in tumor microenvironments engage with Siglecs in order to suppress DC activation and function (73, 74). A similar strategy is also observed in some bacterial infections (75). Sialylated bacterial pathogens like *Campylobacter jejuni* utlize sialylated lipooligosaccharides (LOS) to engage with either siglecs-1 or 7, thereby modulating DC-mediated T cell polarization (75). In a similar scenario, pseudaminic acid-modified flagellin from *Campylobacter jejuni* interacts with Siglec-10 and mediates the secretion of anti-inflammatory cytokine – IL-10 by DCs (15). The sialic acid-Siglec based modulation of DCs has been also studied using artificially produced sialoglycotopes that mimic sialylated products seen in the tumor microenvironment. These sialoglycotopes interact with Siglec-9 to repress IL-12 production by LPS stimulated DCs (32). Artificially sialylated model antigens-ovalbumin (OVA) and myelin oligodendrocyte glycoprotein (MOG_35–55_) interact with Siglec-E on DCs leading to the generation of more Treg cells rather than effector (CD4+/CD8+) T cells (47). In fact, the immunomodulatory role of Siglec-sialoglycans and SHP1/2 phosphatases in cancer is being recognized and targeted to improve anticancer therapy (73, 76). Here, we have observed that the bacterial pathogen *Pseudomonas aeruginosa* is similarly exploiting sialic acid-Siglec based interactions to modulate the process of DC maturation. The DC maturation process is altered by PA Fla^+Sia^ through additional interactions between flagellin sialic acids and the DC surface Siglec-9 receptor. Interestingly, PA is known to switch off the expression of flagellin during chronic infections in order to prevent detection and generation of immune responses against the highly antigenic flagellin (77). In this respect, sialylation of flagellin may be another way by which PA prevents strong immune responses against flagellin. As seen in this current study, immature DCs encountering sialylated PA flagellin leads to Siglec-9 activation and immunosuppression. The resulting signaling cascade also affects the DC maturation process so that these DCs express fewer costimulatory molecules and roduce lowered pro-inflammtory cytokines. More, importantly such interactions also affect the T cell polarization ability of DCs as these DCs generate more Th and Treg cells. The lack of a robust pro-inflammatory response and the production of more Treg cells should be beneficial for PA survival and the establishment of infection.

## Conclusion

DCs play a central role in the immune system as it captures, processes antigens while activating B and T lymphocytes, which form the humoral immunity in the body. The addition of the sialic acid residues gives PA flagellins some new properties. When used as a maturing stimulus, this Fla^+Sia^ interact with both TLR5 and Siglec-9 on immature DC surfaces. The initiation of Siglec based immunoinhibitory signaling affects the TLR5 axis leading to the generation of mature DCs which promote less robust pro-inflammatory responses and produce more Th and Treg cells. Bacterial clearance requires strong but controlled pro-inflammatory responses. Thus, it is likely that PA can utilize silaic acids-Siglec-9 interactions to influence the nature of the maturing DCs as well as regulate the resulting immune response. DCs which promote anti-inflammatory and tolerogenic responses would likely be beneficial for PA survival and dissemination.

## Acknowledgements

KM wants to thank Prof. Chitra Mandal for granting place, funds and resources to conduct this study. KM gratefully acknowledges the encouragements and guidance given by Prof.Chitra Mandal throughout the entire work.

## Supplementary Material

The Supplementary Material for this article is attached at the end of this manuscript.

## Data Availability Statement

All datasets generated for this study are included in the article or the Supplementary Material.

## Author contributions

KM conceived, designed the work with inputs from Prof Mandal.KM performed all the biological experiments and analyzed the data. Manuscript was written by KM with inputs from other senior researchers.

## SUPPLEMENTARY FIGURES

**Figure S1.**
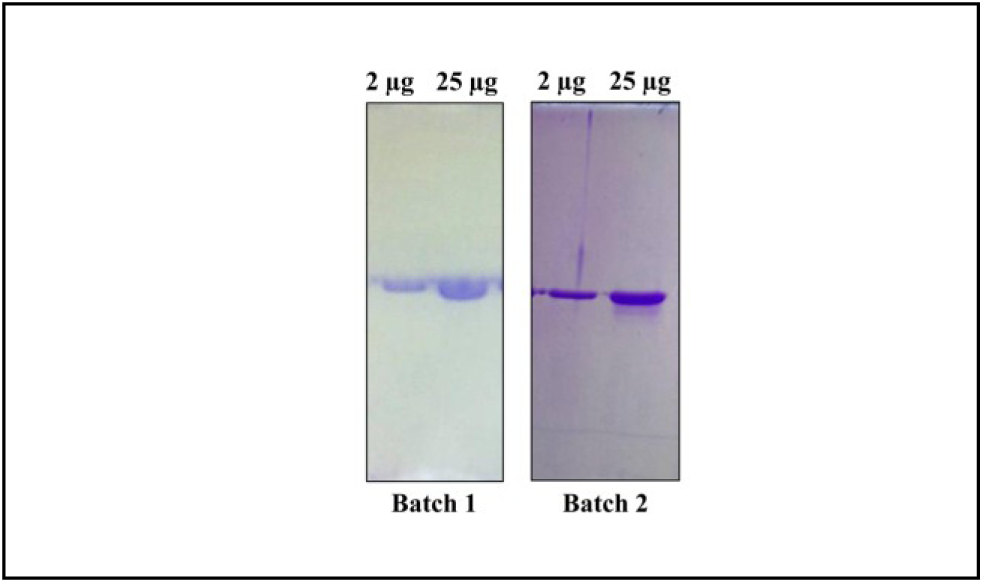
The SDS-PAGE resolution profiles of two different batches of purified Fla^+Sia^ are shown. The presence of single flagellin band even at high amounts of protein loading (25µg) demonstrates the purity of the isolated protein.

**Figure S2.**
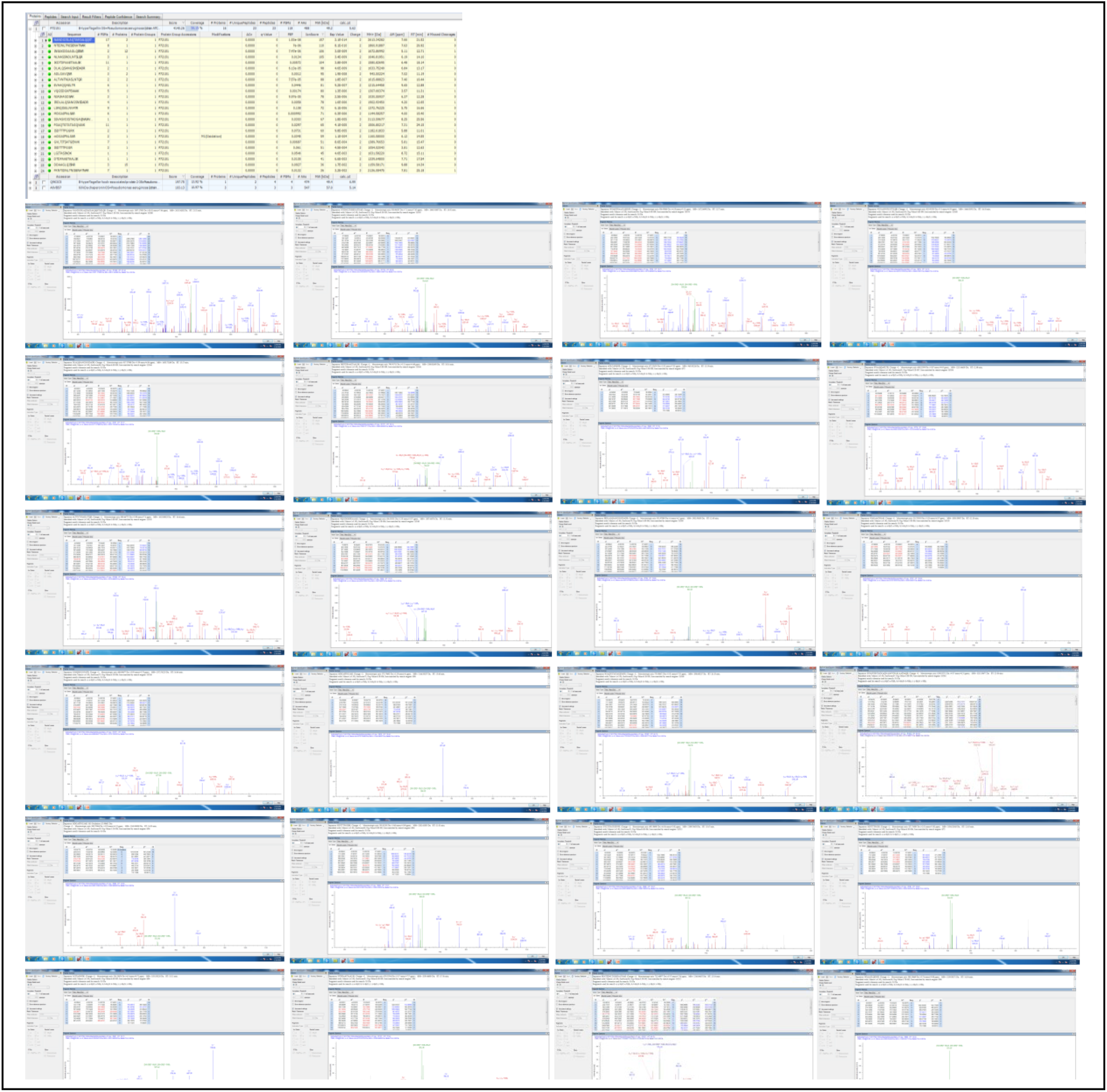
The results of the MASCOT search and the MS/MS spectra generated from various peptide fragments detected in the tryptic digest through nLC-MS/MS (LTQ-Orbitrap) analysis led to the identification of purified Fla^+Sia^.

**Figure S3.**
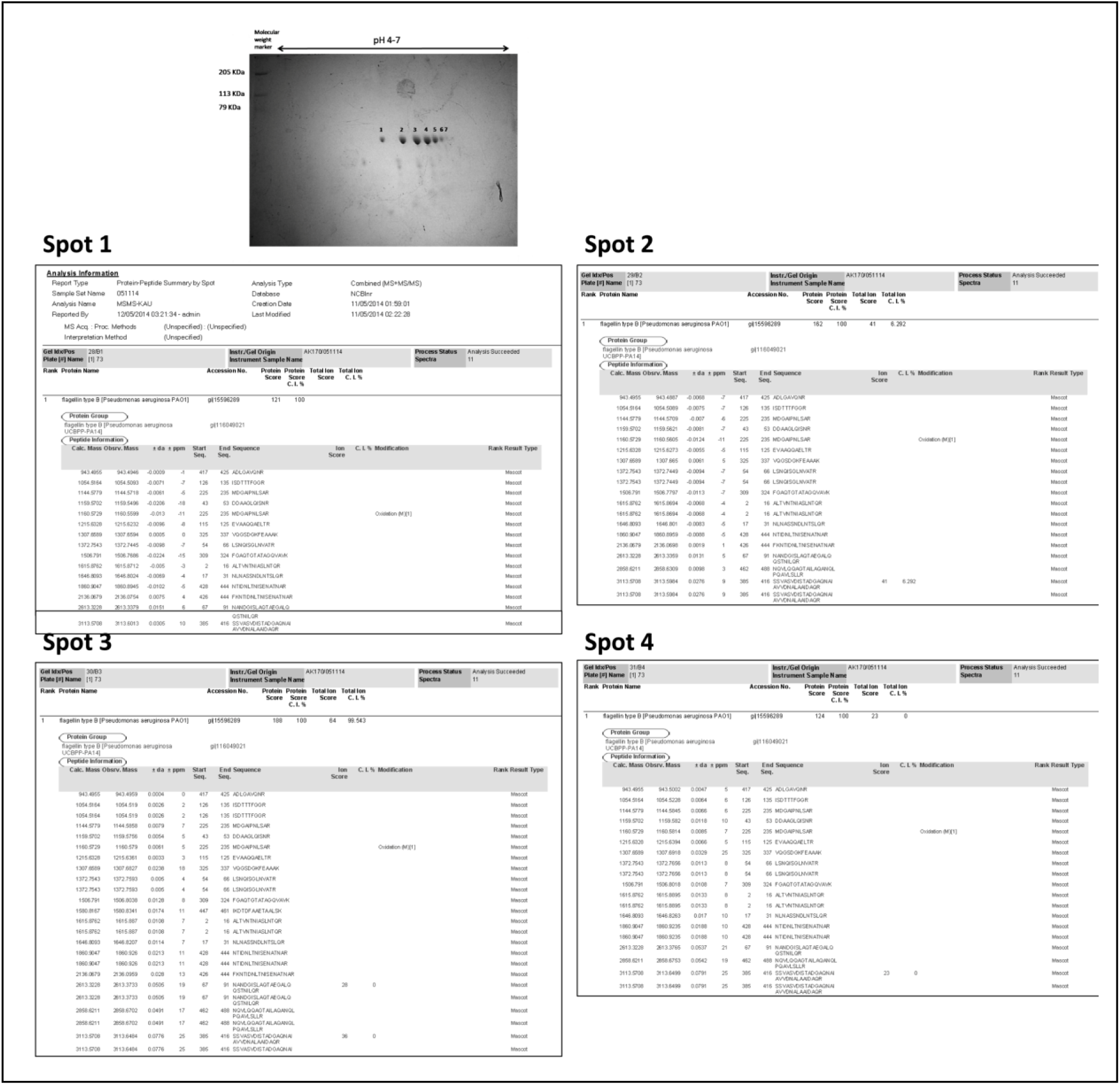
MALDI-TOF MS/MS was used to identify the seven spots observed in 2D electrophoresis of purified Fla^+Sia^ protein. Here, the MASCOT search results of the first 4 spots are shown.

**Figure S4.**
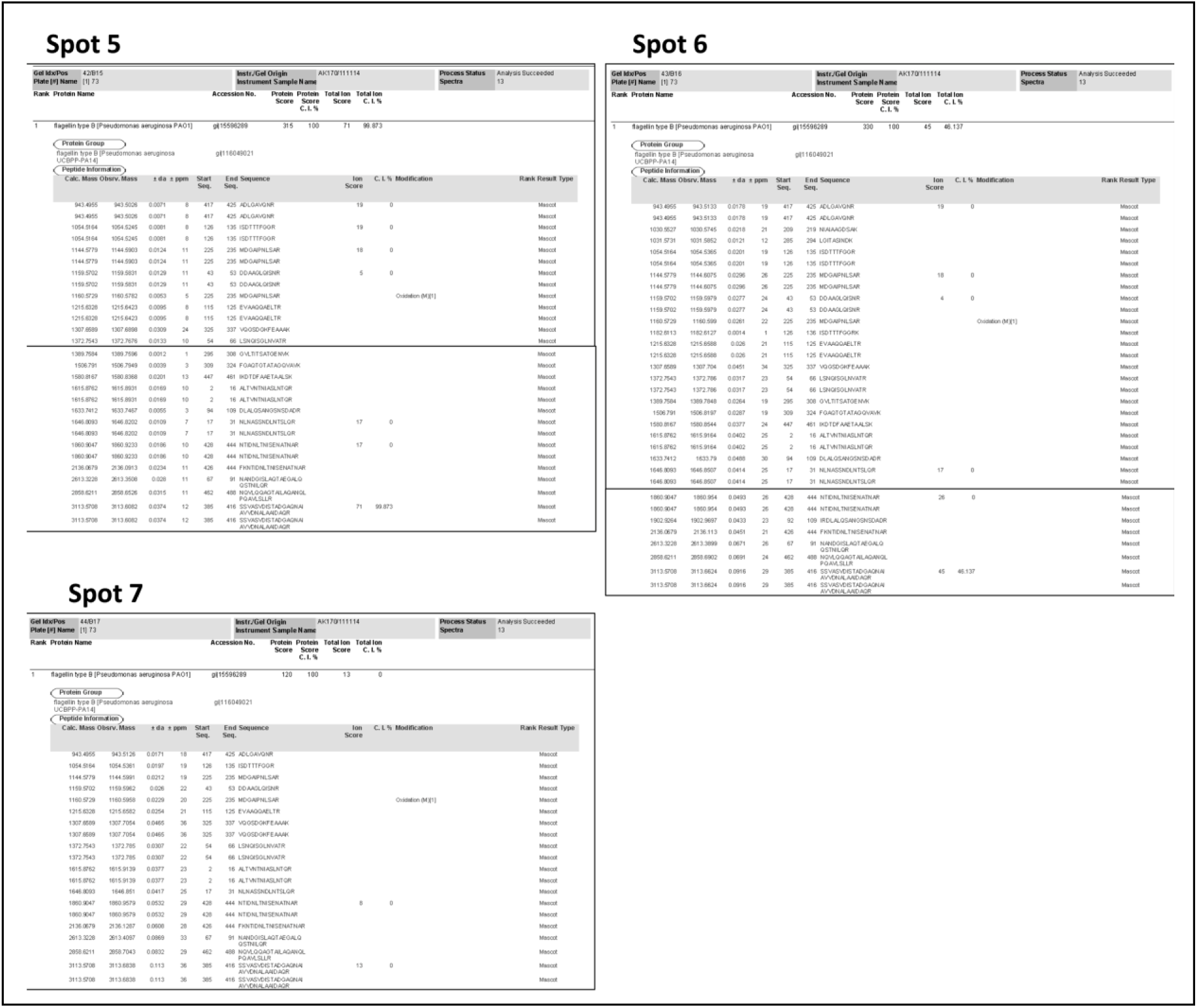
MALDI-TOF MS/MS based identification of the remaining 3 spots out of the seven spots observed in 2D electrophoresis of purified Fla^+Sia^ protein.

**Figure S5.**
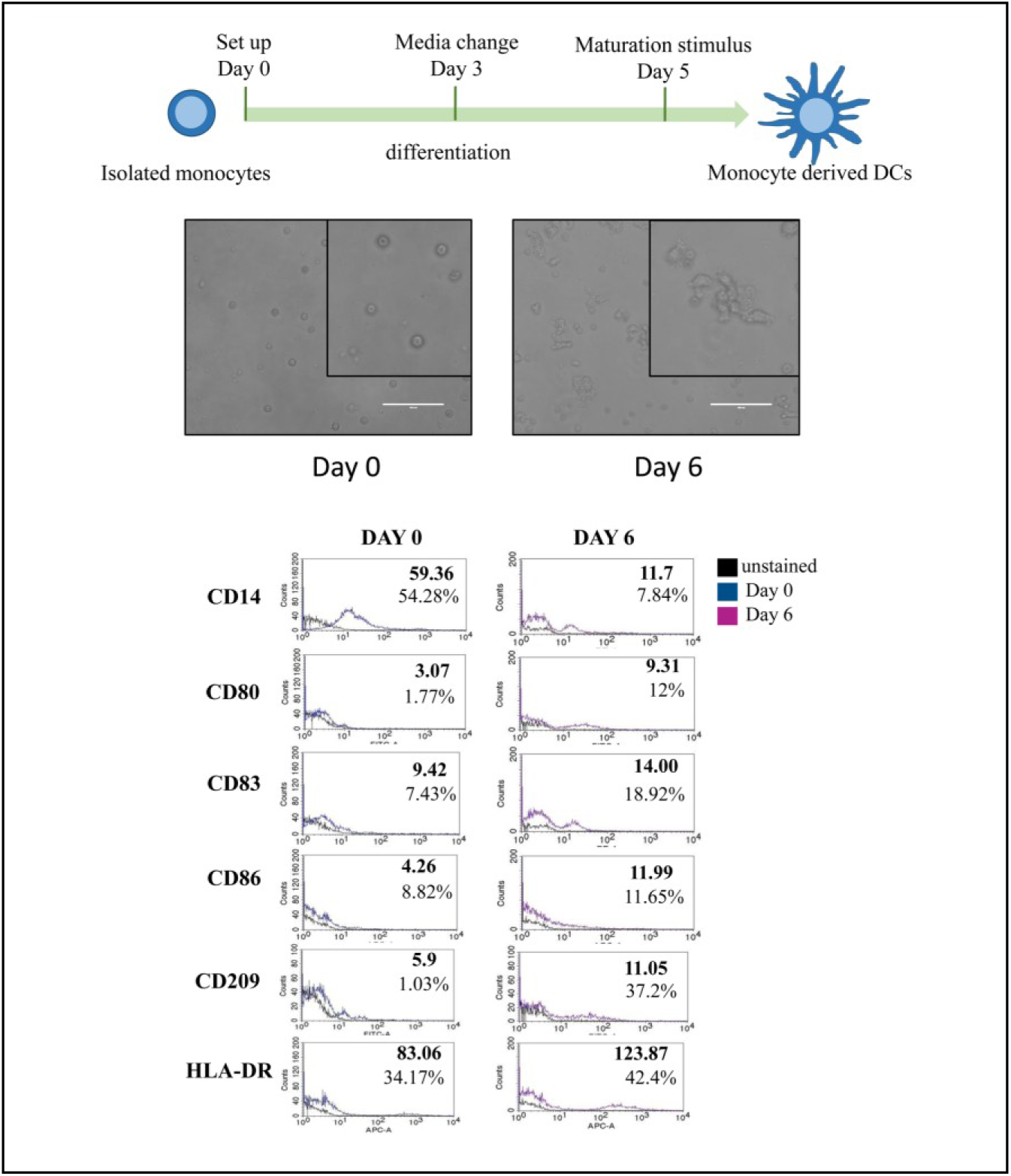
The generation of monocyte derived dendritic cells (moDCs) was monitored by tracking changes in cell morphology and the expression of cell surface receptors.

**Figure S6.**
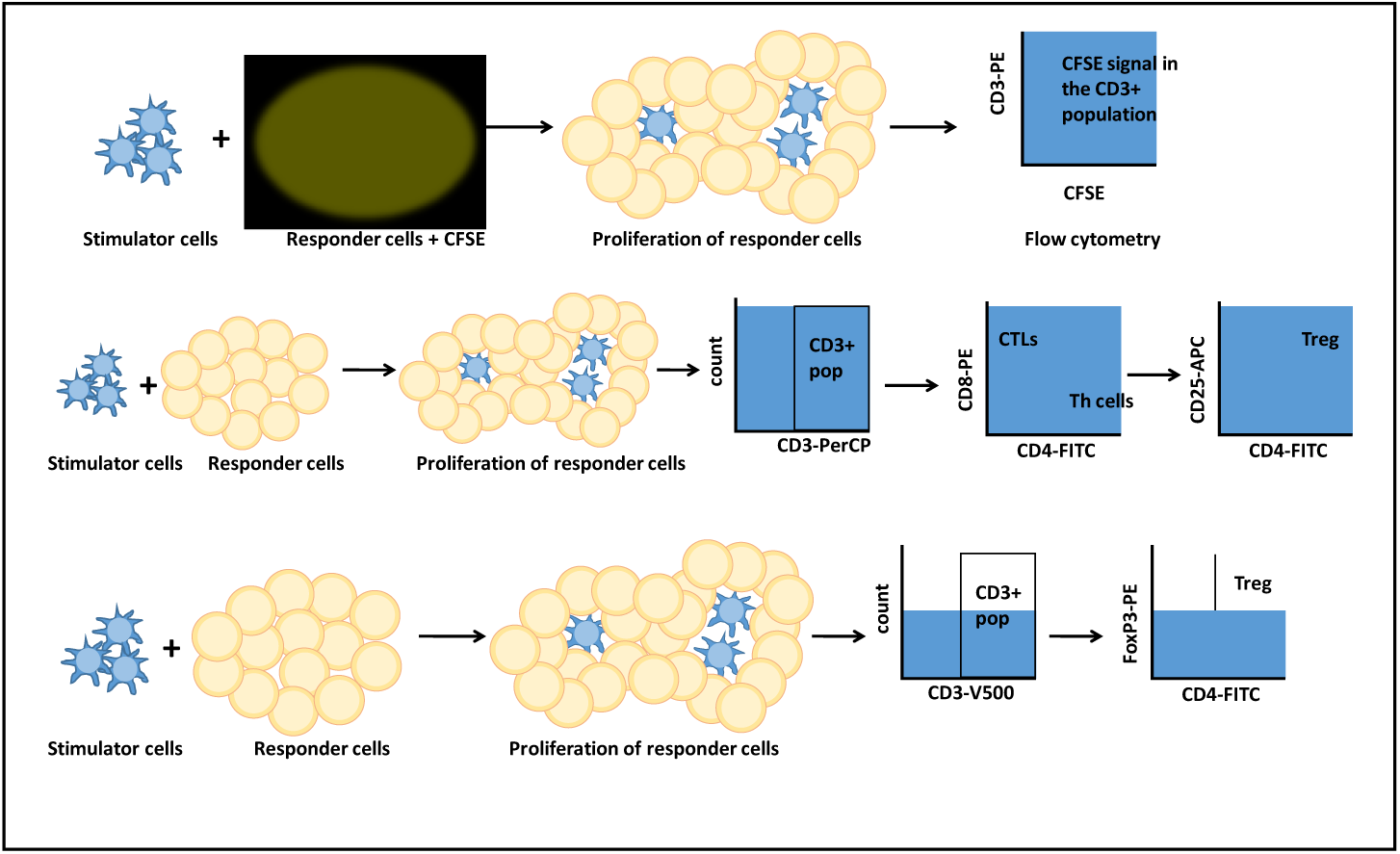
Scheme for the detection of T cell polarization activity by monocyte derived DCs.

**Figure S7.**
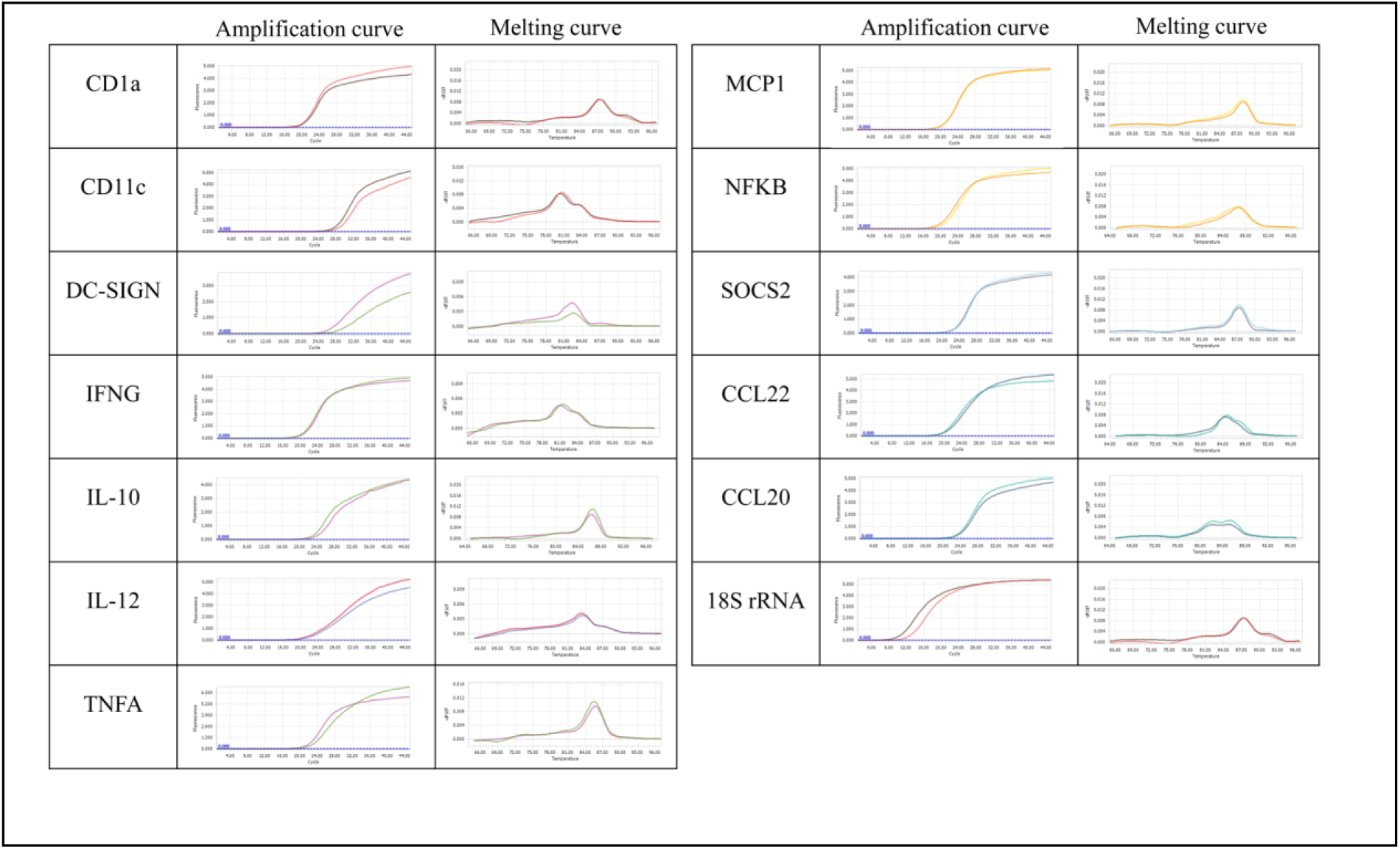
The representative amplification and melting curves of qRT-PCR is shown which is selected from one of the three independent experiments.

